# Higher-order thalamocortical projections selectively control excitability via NMDAR and mGluRI-mediated mechanisms

**DOI:** 10.1101/2023.12.20.572353

**Authors:** Federico Brandalise, Ronan Chéreau, Claudia Morin Raig, Tanika Bawa, Nandkishor Mule, Stéphane Pagès, Foivos Markopoulos, Anthony Holtmaat

**Affiliations:** Department of Basic Neurosciences and the Center for Neuroscience, Centre Médical Universitaire (CMU), University of Geneva, 1211 Geneva, Switzerland; Department of Biosciences, University of Milan, Via Celoria 26, 20133 Milan, Italy; WYSS center, Campus Biotech, Geneva

## Abstract

The apical dendrites of layer (L) 2/3 pyramidal neurons in the mouse somatosensory cortex integrate synaptic input from long-range projections. Among those, inputs from the higher-order thalamic posteromedial nucleus may facilitate sensory-evoked cortical activity, but it remains elusive how this role emerges. Here we show using *ex vivo* dendritic recordings that these projections provide dense synaptic input to broad tufted neurons residing predominantly in L2 and cooperate with other inputs to produce NMDA spikes. They have the unique capacity to block two-pore domain potassium leak channels via group 1 metabotropic glutamate receptor (mGluRI) signaling, which increases excitability. Slender tufted L2/3 neurons and other long-range projections fail to invoke these mechanisms. *In vivo* imaging of calcium signals confirms the presence of mGluRI-dependent modulation of feedback-mediated spiking in L2. Our results imply that higher-order thalamocortical projections regulate neuronal excitability in a cell type and input-selective manner through fast NMDAR and mGluRI-dependent mechanisms.

## INTRODUCTION

The neocortex consists of an intricate network of feedforward and feedback connections, but their topology and functional interactions remain enigmatic. Cortical pyramidal neurons receive distinct synaptic input, depending on their subtype and laminar location^1^. A striking example of spatially segregated inputs to pyramidal neurons is formed by first-order and higher-order parallel thalamocortical projections. In the mouse primary somatosensory cortex (S1), these projections are broadly yet distinctly distributed over all cortical layers^2^. First-order thalamocortical projections from the ventroposterior medial (VPM) thalamus target layer (L) 4 neurons and basal dendrites of thick-tufted L5b and L3 pyramidal neurons, whereas higher-order thalamocortical projections from the posteromedial nucleus (POm) mainly project to L5 and L1, targeting basal dendrites and apical tufts of L5a as well as the apical dendrites of L2/3 pyramidal neurons^3–6^. Within the L2/3 population, these inputs might be biased to L2 neurons^6,7^, but it remains unclear which factors determine this connectivity.

L2/3 pyramidal neurons comprise a morphologically heterogeneous population, with neurons in L2 often bearing extensive apical dendritic tufts, known as broad tufted (BT) neurons, and those in L3 with small tufts, known as slender tufted (ST) neurons^8,9^. However, in mice, the laminar position of pyramidal neurons does not strictly correlate with tuft complexity^10^. The different arrangements of L2/3 pyramidal neuron dendrites are likely to translate into differences in their connectivity, which typically correlates with the amount of axo-dendritic overlap formulated as Peters’ rule^11,12^. Therefore, L2 BT neurons may receive more inputs from long-range axons in L1, whereas L3 ST neurons with disproportionally more basal dendrites may receive biased input from local axons and those terminating in L4 and L3^13–15^. Accordingly, some L2 neurons have been shown to receive relatively strong input from POm thalamocortical projections^7^, although this may not be a general principle for all L2 neurons^6^. Peters’ rule does not apply to all cortical networks. For example, intracortical connectivity of L2 pyramidal neurons is sometimes higher than predicted by axo-dendritic overlap, whereas the input from POm to L5b pyramidal neuron apical tufts in L1 is lower than expected^4,16^. Therefore, it remains unclear if axo-dendritic overlap is a good indicator for the input that L2/3 pyramidal neurons receive from POm afferents.

The connectivity patterns of POm projections suggest that they have distinct roles in the cortical circuitry. This is supported by the notion that synaptic responses evoked by higher-order thalamocortical projections such as from POm, have signatures that are different from synaptic responses elicited by first-order thalamocortical or corticocortical projections^17,18^.

Glutamatergic pathways can be categorized into two groups, termed “drivers” and “modulators.” Driver pathways, such as the pathway from VPM to S1, are linked to information-bearing pathways, whereas modulator pathways, such as the pathway from POm to S1, modify these primary information streams^19,20^. One distinction pertains to the presence of a metabotropic glutamate receptor (mGluR) component^21,22^, but it remains enigmatic how this affects synaptic integration in L2/3 pyramidal neurons. It has been proposed that POm facilitates sensory-evoked responses of pyramidal neurons subpopulations by eliciting long-lasting depolarizations^23–28^, but their underlying mechanisms also remain largely unknown.

Here, by combining electrophysiological dendritic patch-clamp recordings and optogenetics we show that L2/3 neurons with morphologically different dendritic trees receive biased inputs from long-range and local corticocortical circuits. POm thalamocortical synaptic inputs are dense on L2/3 BT neurons, whereas VPM thalamocortical synapses are biased to L2/3 ST neurons. BT neurons produce N-methyl-D-aspartate (NMDA) spikes when POm thalamocortical afferents are stimulated together with other afferents. In addition, we found that POm thalamocortical inputs are unique in their ability to elicit plateau potentials in BT neurons. This effect is mediated by the activation of group 1 mGluRs (mGluRI), which through an interaction with two-pore domain potassium (K2P) leak channels increase the local membrane input resistance. Using 2-photon laser scanning microscopy of calcium signals *in vivo*, we confirm that movement-related activity in these neurons, which is associated with recruitment of feedback circuits, is modulated through mGluRI-mediated mechanisms. We propose that higher-order thalamocortical projections regulate cortical sensory processing by gating the excitability of subpopulations of pyramidal neurons through fast and reversible NMDA receptor (NMDAR) and mGluRI-dependent mechanisms.

## RESULTS

### Distinct long-range inputs to morphological subtypes of L2/3 pyramidal neurons

To compare the relative net input provided by various long-range and local afferents on putatively different types of L2/3 pyramidal neurons, we expressed genetically encoded opsins (ChR2 or ChrimsonR) in putative synaptic afferents using adeno-associated viral (AAV) vectors and recorded from L2/3 pyramidal neuron dendrites in brain slices. During the recordings, cells were filled with biocytin which allowed us to completely reconstruct the morphology of 27 cells for further analysis. To determine the spatial organization of the L2/3 pyramidal neuron dendrites in L1 (Figure 1A and Figure S1) we measured the dendritic density and the span of the tree within the most superficial 200 µm of the somatosensory cortex. Using *k*-means clustering, the neurons were segregated into two groups (Figure 1B), one with reduced and narrow dendritic trees, and one with dense and laterally spreading dendritic trees in L1. The first group typically exhibited a main apical branch extending perpendicular to the pia (Figure S1A), bearing similarities to the formerly reported ST neurons. The second group had dendrites that often originated from two main branches extending in an oblique way towards the pia, similar to the so-called BT neurons^8,14,29^ (Figure S1A). In accordance, both the total length and the number of branches of apical dendrites was larger in BT than in ST neurons whereas the number of branches of basal dendrites was larger in ST than in BT neurons (Figure S1B,C). This resulted in a greatly different apical-to-basal ratio of both branch length and number between the two types (Figure S1B,C), which subsequently allowed us to classify neurons as BT or ST without extensive quantitative reconstructions. In addition, although individual neurons could not be classified as BT or ST neurons based on their laminar position (Figure S1A), the BT neurons were on average located more superficially as compared to ST neurons (Figure 1E), in accordance with previous observations^14^. Pia-aligned dendritic density heatmaps of the two groups indicate that at the population level BT neurons have an overall higher density of dendrites in the L1-L2 region of cortex, whereas ST neurons have slightly more dendritic material in L3 (Figure 1F, see also^30^). Passive electrophysiological parameters of BT and ST neurons were comparable, apart from the slow time constant and capacitance, which was likely a consequence of the morphological differences (Figure S1D,E).

**Figure 1.**
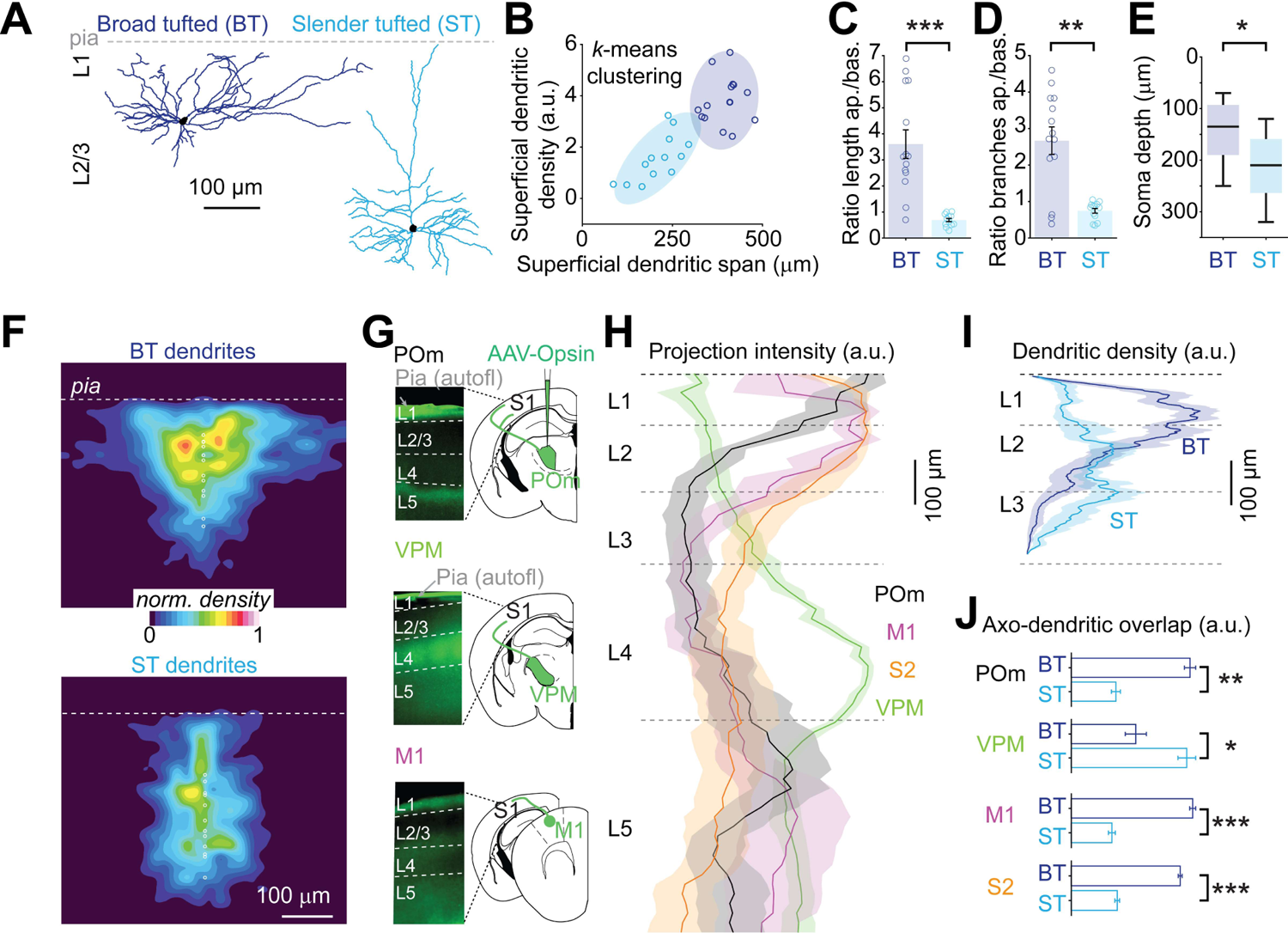
BT and ST neurons in S1 form distinct groups based on morphological features (A) Examples of morphological reconstruction from biocytin-filled L2/3 pyramidal neurons in S1 indicating at least two distinct groups of L2/3 pyramidal neurons as previously described^8,9^: BT neurons exhibiting a large and dense apical arborization and ST neurons displaying a reduced apical arborization. (B) 27 neurons are segregated into BT and ST using a *k*-means clustering method (with *k* = 2) by comparing the dendritic span and density within the first 200 µm from the pia. Ellipsoid areas represent the 95% confidence interval of both clusters. (C) Ratio of the apical over basal dendritic length for BT and ST neurons (*n* = 14 BT neurons and 13 ST neurons, *P* = 5.1×10^-5^, Wilcoxon ranked sum test). (D) Ratio of the apical over basal dendritic branch numbers (*P* = 0.003, Wilcoxon ranked sum test). (E) Comparison of the depth of the soma from the pial surface of BT and ST neurons (*P* = 0.021, Wilcoxon ranked sum test). (F) Dendritic density heatmap of BT and ST neurons aligned to the pia (white circle show somata positions). (G) Examples of long-range projection patterns in S1 after the local injection in POm, VPM and M1 of an AAV vector expressing ChR2-YFP. (H) Fluorescence intensity profiles of POm, VPM, M1 and S2 long-range projections in S1 (each profile is an average from 3 mice and 2 slices per mouse). (I) Dendritic density profiles for BT and ST neurons across cortical layers. (J) Average of the dot product of each long-range input profiles and the dendritic density of each BT and ST neurons (for POm, *P* = 0.002; for VPM, *P* = 0.03; for M1, *P* = 7.7×10^-5^; for S2, *P* = 7.7×10^-5^; Wilcoxon ranked sum test). Error bars, s.e.m.

To estimate the potential synaptic connectivity between long-range thalamocortical or corticocortical afferents and the two types of pyramidal neurons, we compared the laminar distribution of the opsin-labeled axons relative to the reconstructed dendritic trees (Figure 1G,H,I). Thalamocortical afferents from POm and corticocortical afferents from the primary motor cortex (M1) and secondary somatosensory cortex (S2) overlapped more with BT dendrites, whereas thalamocortical afferents from VPM overlapped somewhat more with ST dendrites (Figure 1J).

According to Peters’ rule^11,12^, the patterns of overlap as depicted in Figure 1 predict that BT neurons receive relatively more synaptic inputs from POm, M1 and S2 than ST neurons do, and vice versa, ST neurons should receive relatively more input from VPM in comparison to BT cells. We tested this hypothesis by recording postsynaptic potentials (PSPs) at the main apical dendrite of BT and ST neurons while photo-stimulating the various inputs using 5-ms light pulses (for labeling and recording strategies, see Methods; Figure 2A). Dendritic recordings were used since these are well suited for observing distal dendritic depolarization which readily attenuates toward the soma^31–34^. To account for the variability in the opsin expression levels and patterns over different preparations, we aimed at including both types in each slice to measure the relative synaptic input strength (Figure S2A). Together, this allowed comparisons of input strength from a particular afferent between nearby BT and ST neurons that were surrounded by a similar density of opsin-expressing axons (Figure S2B). The evoked PSP amplitudes increased monotonically with the amount of ChR2-GFP or ChrimsonR-tdTomato fluorescence (Figure S2C). Photo-stimulation of POm afferents evoked PSPs with higher amplitudes in BT neurons as compared to ST neurons (Figure 2B). Conversely, stimulation of VPM afferents evoked larger PSPs in ST neurons as compared to BT neurons. Stimulation of M1 and S2 afferents did not result in statistically different PSP amplitudes between the two types. We also tested the input strength from intracortical S1 inputs (S1_intracortical_) and found no significant differences between the two types (Figure 2B). The PSP rise times were not different between the two types under any of the stimulation conditions (Figure S2D), indicating that the different PSP amplitudes between BT and ST neurons were not due to variations in the distance between the synaptic inputs and the recording sites. To verify that the observed POm and VPM-evoked PSPs included monosynaptic inputs, we bath-applied TTX and 4-AP in a subset of the recordings^4^ (Figure S3A). The application did not abolish POm-evoked PSPs in BT and ST neurons, and VPM-evoked PSPs remained present in ST neurons (Figure S3B). However, the VPM-evoked PSPs in BT neurons were reduced to baseline noise-levels. This data indicates that the responses in ST neurons included monosynaptic PSPs from both POm and VPM, but that BT neurons only receive detectable monosynaptic inputs from POm. Therefore, the VPM-evoked PSPs in BT neurons were likely the result of polysynaptic circuit motifs.

**Figure 2.**
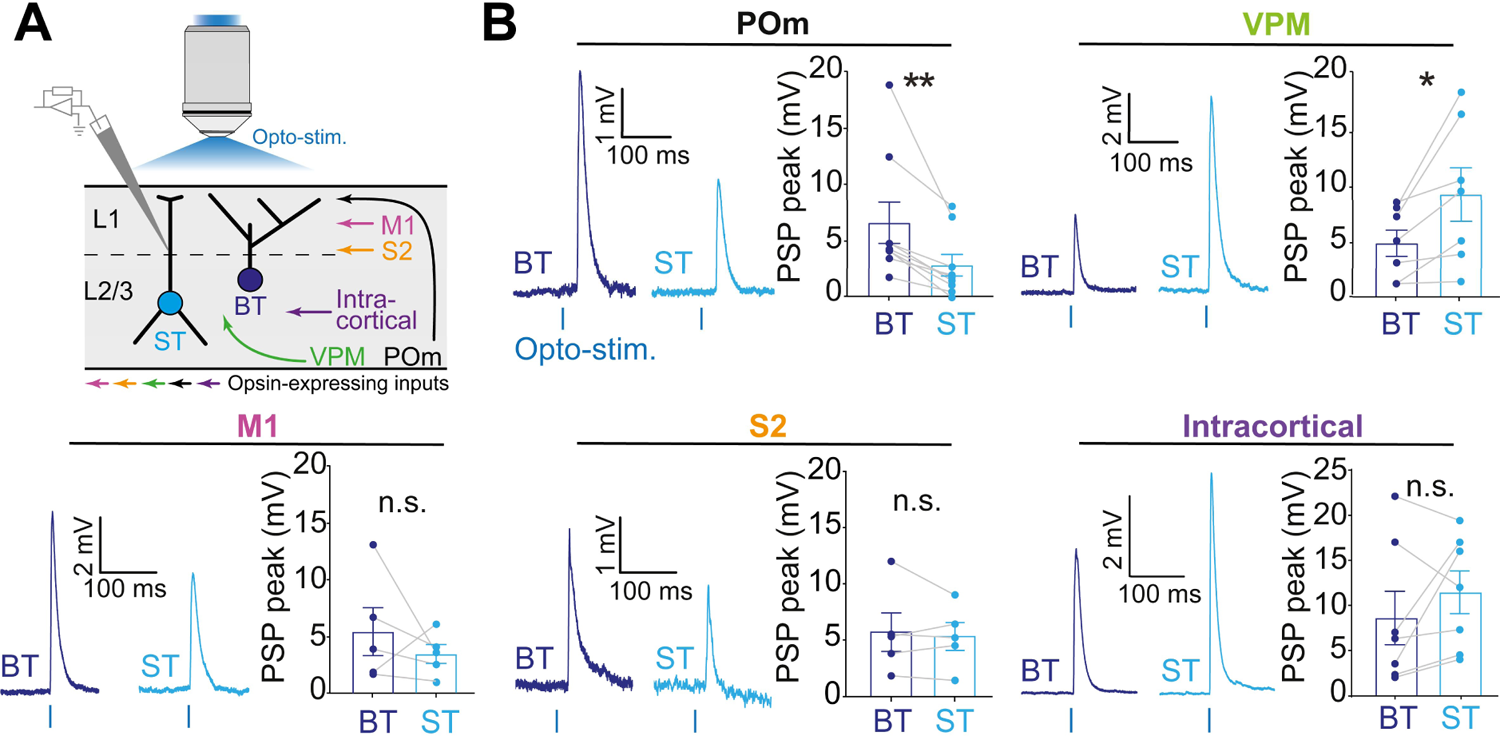
Single afferent input integration on BT and ST neurons in the S1 form distinct groups based on morphological and electrophysiological features (A) Experimental design that consisted in recording the PSP responses of BT and ST neurons upon the photostimulation of various afferent inputs. To account for the variability in expression level of the opsin, a paired comparison between a BT and a ST neuron was performed within each brain slice. (B) Example traces are shown on the left and averaged PSP peaks were calculated for each recorded pairs on the right. BT neurons exhibited larger responses than ST neurons when POm was stimulated (*n* = 9 pairs; BT: 6.5 ± 1.8 mV; ST: 2.8 ± 0.9 mV; *P* = 0.004, Wilcoxon signed-rank test). However, ST neurons had significantly larger responses than BT neurons when VPM was stimulated (*n* = 7 pairs; BT: 4.8 ± 1.2 mV; ST: 9.3 ± 2.4 mV; *P* = 0.0154, Wilcoxon signed-rank test). No significant differences were observed when M1, S2 and S1_intracortical_ inputs were stimulated (for M1, *n* = 5 pairs; BT: 5.3 ± 2.3 mV; ST: 3.4 ± 3.7 mV; *P* = 0.43; for S2, *n* = 5 pairs; BT: 5.7 ± 1.7 mV; ST: 5.3 ± 1.2 mV; *P* = 0.94, for S1_intracortical_, *n* = 7 pairs; BT: 7.7 ± 3.3 mV; ST: 12.0 ± 3.4 mV; *P* = 0.2 Wilcoxon signed-rank tests). Error bars, s.e.m.

To further investigate the different levels of thalamocortical input to the two groups of neurons, we utilized the anterograde trans-synaptic labeling properties of AAV1^35^. AAV1-mCaMKIIα-iCre-WPRE-hGHp(A) was injected in either the POm or VPM, and AAV2-hSyn-DIO-eGFP in S1 (Figure S4A). Owing to the trans-synaptic transport of the AAV1-Cre vector, GFP expression was driven in the neurons that held synaptic connections with thalamocortical axons. AAV1-Cre from the POm drove GFP expression predominantly in cortical L2/3 and L5 neurons, whereas from the VPM it labeled cells in L4 and L2/3 (Figure S4B). The L2/3 neurons that were targeted by POm afferents were on average located closer to the pial surface than those targeted by VPM afferents (Figure S4C). Since BT neurons tend to be positioned more superficially in the cortex as compared to ST neurons (Figure 1E), these observations suggest that POm axons target predominantly BT neurons, whereas VPM axons are biased to ST neurons. This corroborates our electrophysiological findings showing that BT neurons received on average stronger synaptic input from POm, and ST neurons stronger input from VPM (Figure 2B).

### Inputs from POm afferents combined with other long-range synaptic inputs selectively induce NMDAR-dependent responses in BT neurons

Cortical L2/3 pyramidal neurons can integrate dendritic inputs in a supralinear manner, mediated by NMDARs, also called NMDA spikes^23,33,36–39^. These events are facilitated under depolarized conditions, when synapses are clustered, or when synapses harbor signaling mechanisms that strongly interact with one another^40–43^. L2/3 neurons have been shown to produce NMDA spikes upon sensory stimulation which may depend on inputs from POm and other afferents in L1^23,39^. Therefore, we sought to investigate whether the stimulation of different combinations of thalamocortical and corticocortical afferents have distinct propensities to produce NMDA spikes in BT and ST neurons. We expressed ChR2 and ChrimsonR in various pairs of putative presynaptic afferents and then performed dendritic recordings from either class in brain slices while photostimulating two afferents simultaneously (Figure 3A). We used light intensities and wavelengths that generated action potentials in the opsin-expressing neurons, but avoided cross-contamination between the two light channels (Figure S5, see^44^). The recorded dendrites were held at −55 mV to facilitate the generation of NMDA spikes. We first simultaneously photostimulated POm and M1 afferents. In BT cells this evoked seemingly two types of PSPs, characterized by smaller and larger amplitudes (Figure 3B). The large-amplitude PSPs were prevented upon perfusion of the NMDAR antagonist APV, but the smaller amplitude PSPs remained unaffected. The large-amplitude PSPs were also prevented when NMDAR opening was precluded by holding cells at hyperpolarized potentials (∼-100mV) (Figure S6A). Together, this confirms that the larger PSPs included NMDAR-mediated conductance and can be classified as NMDA spikes (Figure 3B). NMDA spikes were also observed in ST neurons upon stimulation of VPM and S1_intracortical_ afferents (Figure S6B). To assess the efficacy by which various afferent pairs evoked NMDA spikes, we first determined whether the distribution of the evoked PSP amplitudes was bimodal (see methods, Figure 3B, and Figure S6). Then, for each combination of inputs and each cell type, *k*-means clustering was used to separate the NMDA spikes from the regular PSPs, from which the fraction of trials with NMDA spikes as well as their total strength (fraction multiplied by amplitude) were computed. This analysis revealed that co-stimulation of POm and M1 afferents had a higher capacity to evoke NMDA spikes in BT neurons as compared to ST neurons, as well as compared to the other tested combinations of putative inputs (Figure 3C). Most BT neurons also displayed NMDA spikes when POm and VPM afferents were co-stimulated. This is intriguing since we could not detect distinct monosynaptic inputs coming from VPM afferents onto BT neurons (Figure S3). It implies that the stimulation of POm afferents combined with VPM-mediated activation of local excitatory circuits such as from L4 and ST neurons, can generate NMDA spikes. The generation of NMDA spikes in ST neurons was most pronounced when VPM afferents and S1_intracortical_ circuits were co-stimulated (Figure 3C). These data indicate that NMDA spikes can be evoked in L2/3 pyramidal neurons upon combined stimulation of long-range synaptic input. This bears similarities to the NMDA spikes that have been reported in L4 upon combined thalamocortical and intracortical stimulation^38^, and the supralinear potentials in L2/3 pyramidal neurons upon sensory stimulation^23,39^. Our data show that these NMDA spikes are L2/3 neuron type and input selective.

**Figure 3.**
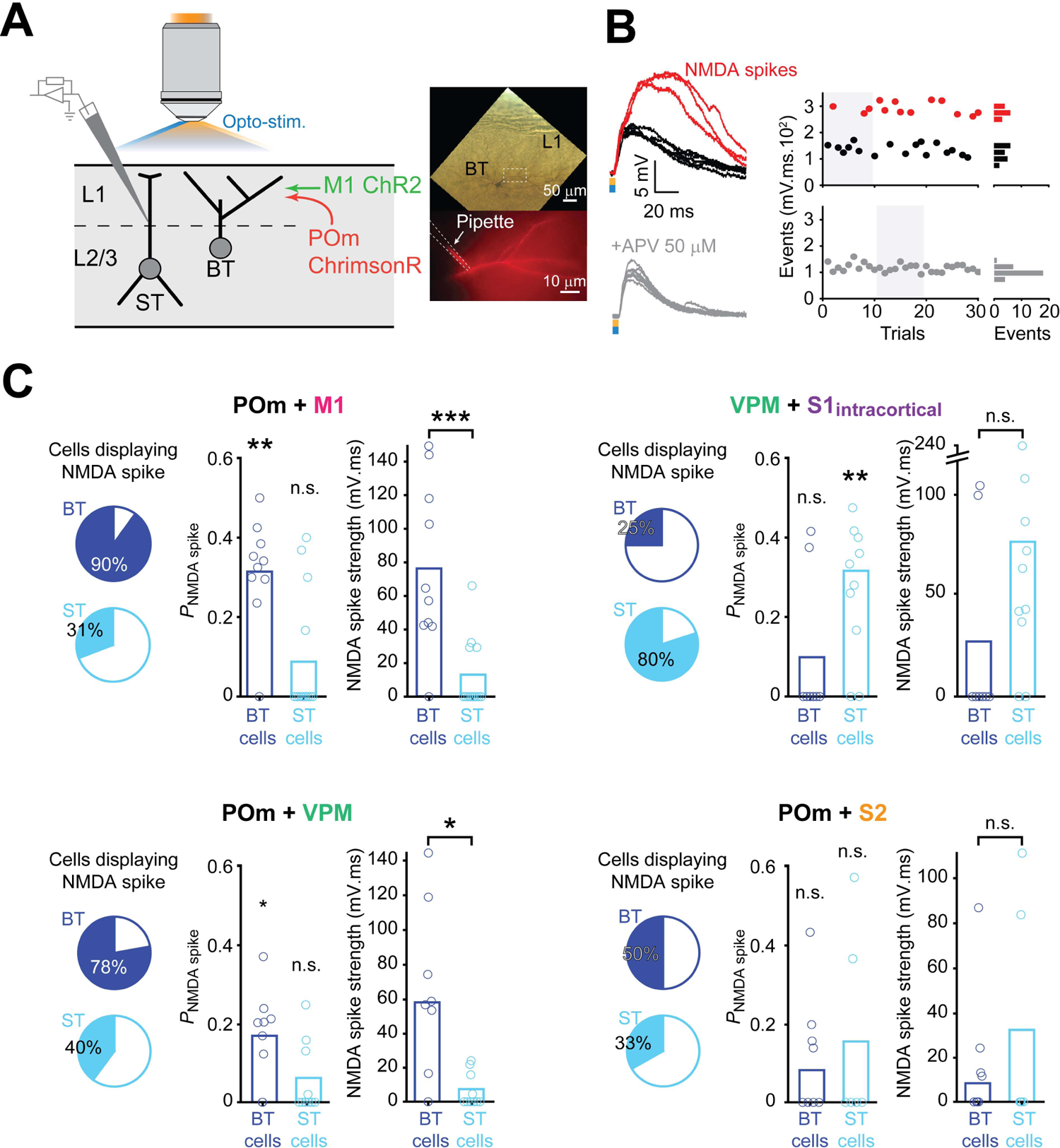
Cell-type and input specific generation of NMDA spikes in L2/3 neurons (A) Experimental design for testing the integration of two inputs that converge onto BT and ST neurons in S1. The two inputs expressed different opsins (ChR2 and ChrimsonR) by local injection of AAV vectors. In this example, ChrimsonR was expressed in POm and ChR2 in M1. To ensure accurate recording of distal events, patch-clamp recordings were performed on the apical dendrites of BT or ST neurons. (B) Examples of events evoked by the co-stimulation of POm and M1 inputs on a BT neuron (top). The stimulation either elicited regular PSPs (black traces) or NMDA spikes (red traces). The bath application of 50 mM APV prevented the generation of NMDA spikes (bottom). The distribution of the size of the events, from this example, showing that NMDA spikes can be easily segregated from regular PSPs (right). (C) Occurrence of NMDA spikes for four different input combinations. For each of them, the pie charts indicate the percentage of BT and ST cells that displayed NMDA spikes at least once during the recording period (left). For each of the cell types, the NMDA spike probability per trial was compared to the null hypothesis of a zero probability (middle, for POm + M1: BT cells, *n* = 10, *P* = 3.9×10^-3^; ST cells, *n* = 13, *P* = 0.12; for VPM + S1_intracortical_: BT cells, *n* = 8, *P* = 0.5; ST cells, *n* = 10, *P* = 7.8×10^-3^; for POm + VPM: BT cells, *n* = 9, *P* = 0.02; ST cells, *n* = 10, *P* = 0.12; for POm + S2: BT cells, *n* = 8, *P* = 0.12; ST cells, *n* = 6, *P* = 0.5; Wilcoxon rank sum tests). The NMDA spike strength between BT and ST groups was compared (right, for POm + M1: *P* = 8.7×10^-3^; for VPM + S1_intracortical_: *P* = 0.09; for POm + VPM: BT cells, *P* = 0.054; for POm + S2: BT cells, *P* = 0.93; same ns as for the fraction of trials, Wilcoxon sign rank tests).

### Inputs from POm afferents evoke plateau potentials in BT cell dendrites

While we assessed the functional connectivity of long-range inputs onto L2/3 pyramidal neurons, we observed that optogenetic stimulation of POm afferents using a train of 5 pulses (8Hz) evoked sustained plateau-like depolarizations that followed the 5 short-latency PSPs with long and variable delays (Figure 4A). Such events were virtually absent upon stimulation of other afferents.

**Figure 4.**
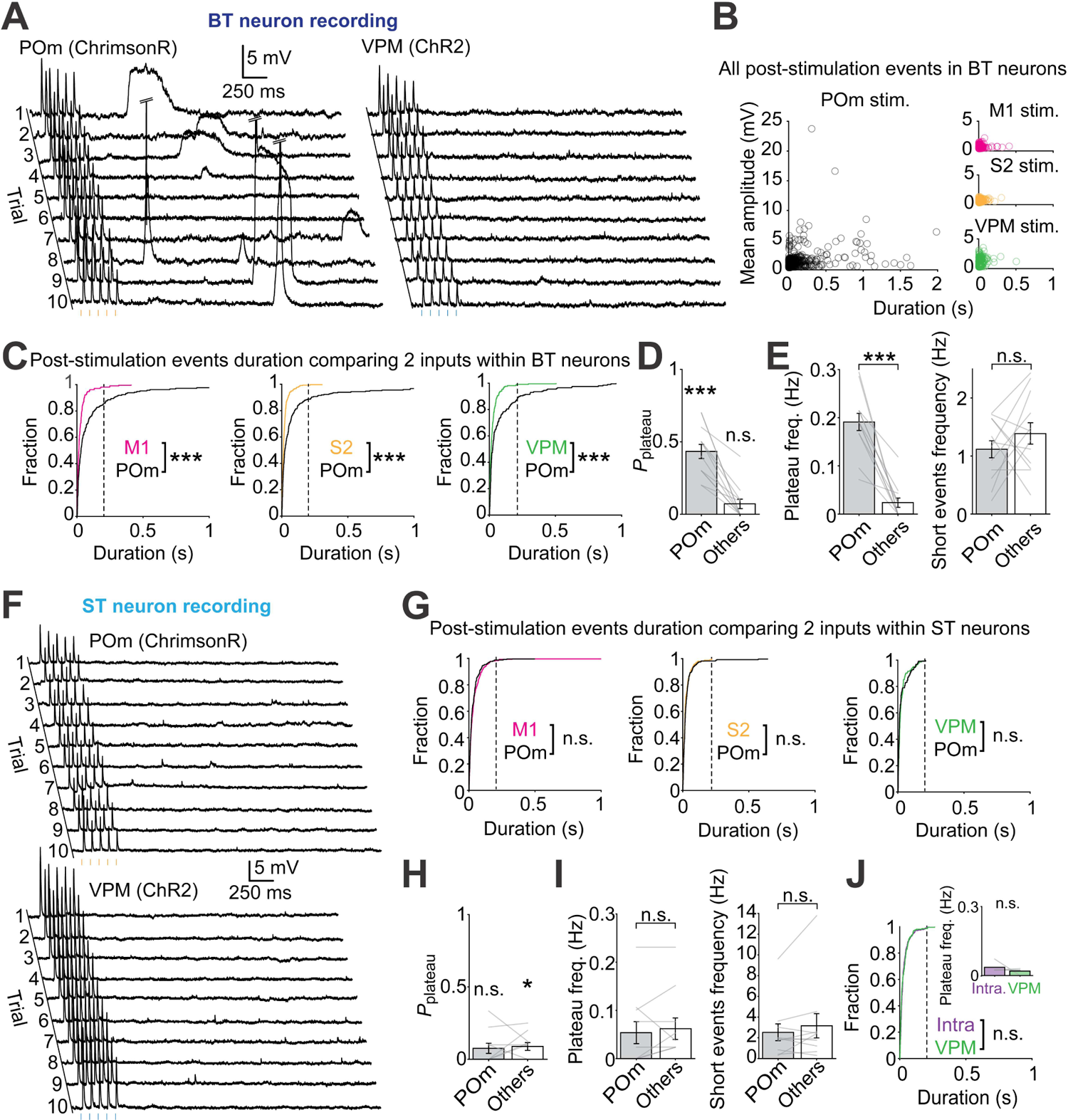
POm activation induces long-lasting and delayed plateau potentials in BT neurons (A) Example traces of the dendritic recording from a BT neuron when POm afferent inputs expressing ChrimsonR and VPM afferent inputs expressing ChR2 were photostimulated independently. In both cases, the five pulses of light elicited PSPs. Long-lasting plateau potentials were regularly observed after POm was stimulated. These events occurred with a highly variable delay and were not observed following VPM stimulation. (B) Scatter plots showing the duration and mean amplitude of all post-stimulation events that were automatically detected in BT neurons. While only short duration events of small amplitude, corresponding to spontaneous PSPs, were detected following the stimulation of M1, S2 or VPM, long duration events of variable amplitude were detected following POm stimulation. (C) Cumulative fractions of the post-stimulation event durations for POm and another input (M1, S2 and VPM) in BT neurons (left, M1 vs. POm: *P* = 1.7×10^-7^, S2 vs. POm: *P* = 1.6×10^-10^, VPM vs. POm: *P* = 5.5×10^-5^, Koglomorov-Smirnov tests). (D) Probability of detecting at least one plateau potential following optogenetic stimulation of POm and optogenetic stimulation of other M1, S2 and VPM inputs tested against the zero probability (*n* = 12, for POm: *P* = 4.8×10^-4^; for others: *P* = 0.06, Wilcoxon sign-rank tests). (E) Left, plateau potentials frequency in BT neurons for POm compared to other inputs (*n* = 12, *P* = 4.8×10^-4^; Wilcoxon sign-rank test). Right, same comparison for short events frequency (*P* = 0.37; Wilcoxon sign-rank test). (F) Example traces of the dendritic recording of an ST neuron after independent photostimulation of POm and VPM afferent inputs. No plateau potentials were observed when POm or VPM were stimulated. (G) Cumulative fractions of the post-stimulation event durations for various pairs of inputs in ST neurons (left, M1 vs. POm: *P* = 0.58, S2 vs. POm: *P* = 0.35, S1_intracortical_ vs. VPM: *P* = 0.055, VPM vs. POm: *P* = 0.052, Koglomorov-Smirnov tests). (H) Same analysis as (D) for ST neurons (*n* = 10, for POm, *P* = 0.12; for other inputs: *P* = 0.03; Wilcoxon sign-rank test). (I) Same analysis as (E) for ST neurons (*n* = 10, for plateau potential frequency, *P* = 0.62; for short events frequency, *P* = 0.32, Wilcoxon sign-rank test). (J) Cumulative fractions of the post-stimulation event durations for S1_intracortical_ and VPM inputs in ST neurons (*P* = 0.054, Koglomorov-Smirnov tests). Inset shows the plateau potentials frequency in ST neurons for this pair of inputs (*n* = 4, *P* = 0.44; Wilcoxon sign-rank test).

To better characterize these events, we compared pairs of inputs on BT and ST neurons from ChR2 and ChrimsonR expressing afferents, which were independently tested at least 10 times by interleaving the two optogenetic stimuli every 10 s. Dendritic recordings from BT neurons systematically displayed long-lasting depolarizing potentials following optogenetic stimulation of POm but not of VPM (Figure 4A). We did not observe this in ST neurons (Figure 4F). To quantitatively assess these events, we designed a filter to detect any depolarization that occurred after the 5 stimuli, and then measured their durations and amplitudes. Stimulation of POm afferents often evoked large-amplitude and long-lasting events, which were not seen after stimulation of M1, S2, and VPM (Figure 4B). The cumulative distribution of the event durations indicates that inputs from POm consistently produced longer-lasting events as compared to the other inputs (Figure 4C). A similar analysis of the recordings from ST neurons did not reveal differences in the cumulative distribution of event durations between the POm simulation and any other inputs (Figure 4F,G).

These long-lasting depolarizations were clearly distinct from regular PSPs and NMDA spikes. Their average duration, rise time, and amplitudes were significantly different from the NMDA spikes seen upon combined stimulation of inputs (Figure S7). Together, this indicates that these events were very unlikely to represent spontaneous AMPAR-mediated PSPs or NMDA spikes. To distinguish them from NMDA spikes, we termed them plateau potentials for the remainder of the paper.

The probability of detecting plateau potentials (>200ms) in BT neurons after stimulation of POm afferents was significantly higher than after stimulation of any of the other inputs (Figure 4D). Even though they sporadically arose after stimulation of the other inputs, their probability was not significantly above zero (Figure 4D). Thus, the increased frequency of plateau potentials appeared exclusively associated with stimulation of POm afferents (Figure 4E). The increase in plateau potential frequency was independent of the short duration event frequency, which was not different between POm and other afferent stimulation (Figure 4E). Stimulation of POm afferents did not increase their probability in ST neurons (Figure 4H) as compared to other afferent stimuli, and no difference was found when comparing the frequency of plateau potentials and short-lasting events (Figure 4I). Moreover, when comparing the VPM and S1_intracortical_, we almost exclusively found short-lasting events and the measured frequencies of plateau potentials were close to zero (Figure 4J). In addition, BT neurons in M1 did not produce any plateau potentials upon optogenetic stimulation of POm afferents (Figure S8). Together, these results indicate that plateau potentials predominantly occurred in S1 BT neurons and were selectively associated with the stimulation of POm inputs.

### Plateau potentials depend on the closing of leak K^+^ channels

Since the plateau potentials were so distinct from any of the other events, we hypothesized that they were associated with the opening or closing of ion channels other than the typical synaptic receptors. To investigate the conductance that was mediating the plateau potentials, we recorded from BT neurons while optogenetically stimulating POm afferents and holding the membrane potential at −60, −100 and −125 mV (Figure 5A). Whereas the plateau potentials were depolarizing at −60 mV, they nearly disappeared at −100 mV and became hyperpolarizing at −120 mV. We inferred that the reversal potential of these events was around −104 mV (Figure 5B). This is consistent with a potassium conductance, which under our experimental conditions was estimated to be around −100 mV (see methods). Considering that plateau potentials were detected as depolarizing at resting membrane potentials, we deemed it unlikely that they were mediated by the opening of hyperpolarizing and voltage-dependent potassium channels. Instead, we hypothesized that they were mediated by leak potassium channels that regulate resting membrane potentials, the majority of which is formed by the K2P channel family^45^. Under our experimental conditions, the plateau potentials would thus reflect the transient closing of the K2P channels. To test this, we performed dendritic recordings of BT neurons, while optogenetically stimulating POm afferents. We measured the plateau potential frequency before and after bath application of a broad-spectrum cocktail of K2P channels blockers^46^ (Figure 5C). Consistent with the hypothesis, the blocking of these channels significantly reduced the frequency of plateau potentials (Figure 5C,D). To narrow down which K2P channel subtypes could be involved, we tested the effect of more specific antagonists in a different set of experiments. Blocking TASK or TREK channels by bath application of A1899 or fluoxetine significantly reduced the plateau potentials frequency, whereas the blocking of THIK-1 channels by IBMX did not affect them (Figure 5G). Finally, the blocking of K2P channels by any of these means increased the membrane resistance of the recorded neurons (Figure 5F), suggesting that the effects were cell autonomous.

**Figure 5.**
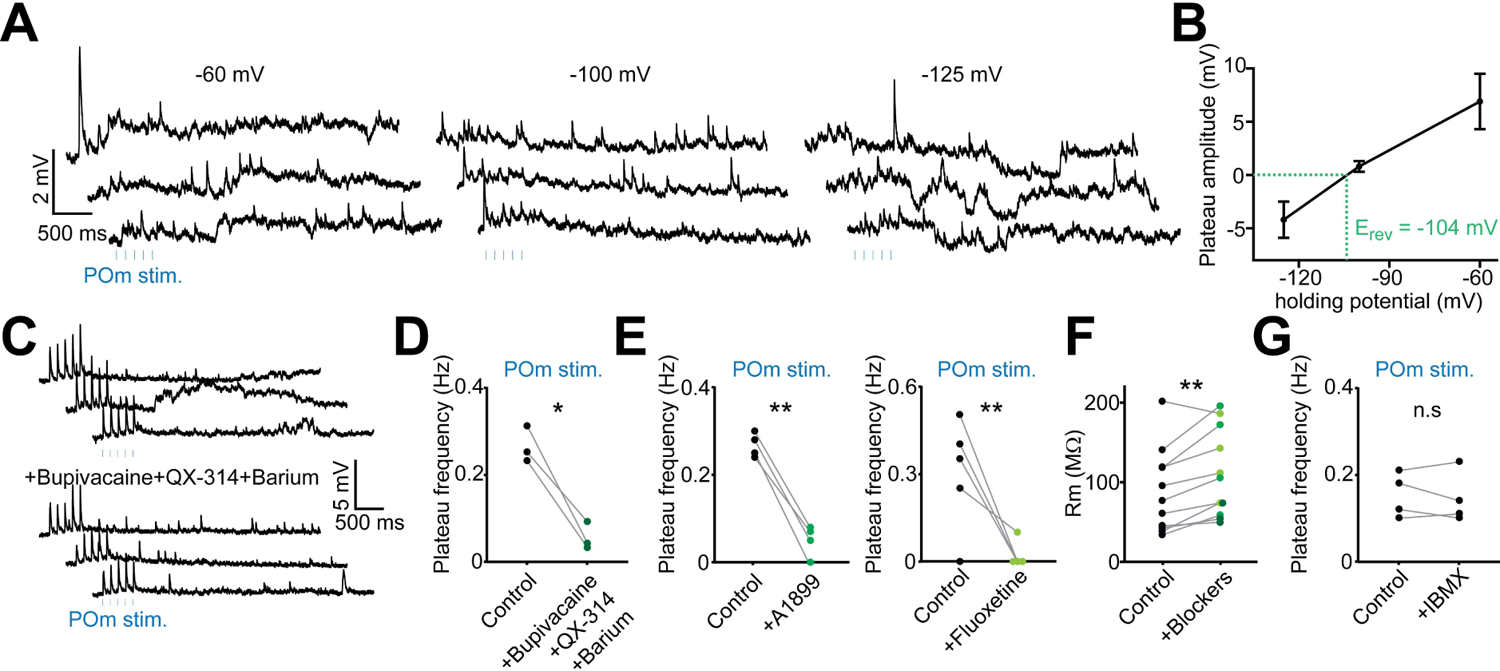
POm-mediated plateau potentials are due to the closing of K2P channels (A) Dendritic recordings of a BT neuron held at different holding potentials during the photostimulation of POm (5 pulses of 1 ms at 8 Hz, every 10 s). Plateau potentials are observed at a holding potential of −60 mV as long-lasting depolarizing events. Applying a holding potential of −125 mV reverted the direction of these events by displaying hyperpolarizing plateau potentials. (B) Plateau amplitude evoked by the stimulation of POm as a function of the holding potential (*n* = 3 neurons). The reversing potential of these events was measured at −104 mV consistent with a potassium conductance (see methods). (C) Dendritic recordings of a BT neuron during the stimulation of POm (5 pulses of 1 ms at 8 Hz, every 10 s) before and after bath application of bupivacaine (1 mM), QX-314 (1 mM) and barium (1 mM). No plateau potentials could be observed in the presence of this non-specific blockade of K2P channels. (D) Plateau potentials frequency was significantly reduced in the presence of non-specific K2P channels blockers (*n* = 3, *P* = 0.023, paired t-test). (E) The selective blockade of TASK or TREK channels using A1899 (100 nM) or fluoxetine (100 µM) respectively, largely prevented the generation of plateau potentials (for A1899, *n* = 4, *P* = 0.001; for fluoxetine, *n* = 5, *P* = 0.005, paired t-tests). (F) Altogether, the K2P blockers used in D and E significantly increase the membrane resistance of the recorded dendrites (*n* = 11, *P* = 0.004, paired t-test). (G) Blocking the THIK channel family with IBMX (1 mM) did not produce any change in the frequency of plateau potentials (*n* = 4, *P* = 0.62, paired t-test).

### Plateau potentials in POm to BT synaptic inputs are mediated by post-synaptic group I mGluRs

We next sought to investigate how the activation of POm to BT synaptic inputs leads to the blocking of TASK/TREK channels. TASK and TREK channels have been shown to be modulated by G protein-coupled receptor (GPCR) signaling pathways^47,48^. The mGluRIs have been shown to induce delayed and long-lasting depolarizing events resembling the ones observed in our recordings^21,22,49^. Therefore, we hypothesized that the activation of POm to BT synaptic inputs triggers mGluRI-signaling which subsequently mediates the transient closing of TASK or TREK channels. To test this, we performed another set of dendritic recordings on BT neurons and measured the frequency of plateau potentials following POm stimulation before and after bath perfusion of specific and generic mGluRI blockers, LY367385 and MCPG respectively (Figure 6A). Blocking mGluRIs significantly reduced the frequency of plateau potentials, similar to the effect of blocking of K2P channels (Figure 6A,B). To confirm that the effects sorted by LY367385 and MCPG were mediated by postsynaptic mGluRI, we added GDP-β-S to the intracellular solution (Figure 6D). GDP-β-S is a non-hydrolyzable analog of GDP that internally blocks G-protein activity^17,50^. Whereas optogenetic stimulation of POm still induced plateau potentials immediately after break-in with the patch electrode, the events were largely abolished within approximately 3 minutes (Figure 6D,E), which is consistent with the dialysis kinetics of GDP-β-S^51^. Overall, the presence of mGluRIs or G-protein blockers slightly decreased the membrane resistance of all recorded neurons (Figure 6C,F). Conversely, the bath application of the mGluRI agonist DHPG increased both the frequency and the amplitude of plateau potentials triggered by POm stimulation, along with an increase in the input resistance (Figure S9A-C). This effect was absent in ST neurons (Figure S9D,E). Taken together, these experiments indicate that plateau potentials are mediated by the signaling of postsynaptic mGluRIs at the POm to BT synaptic inputs.

**Figure 6.**
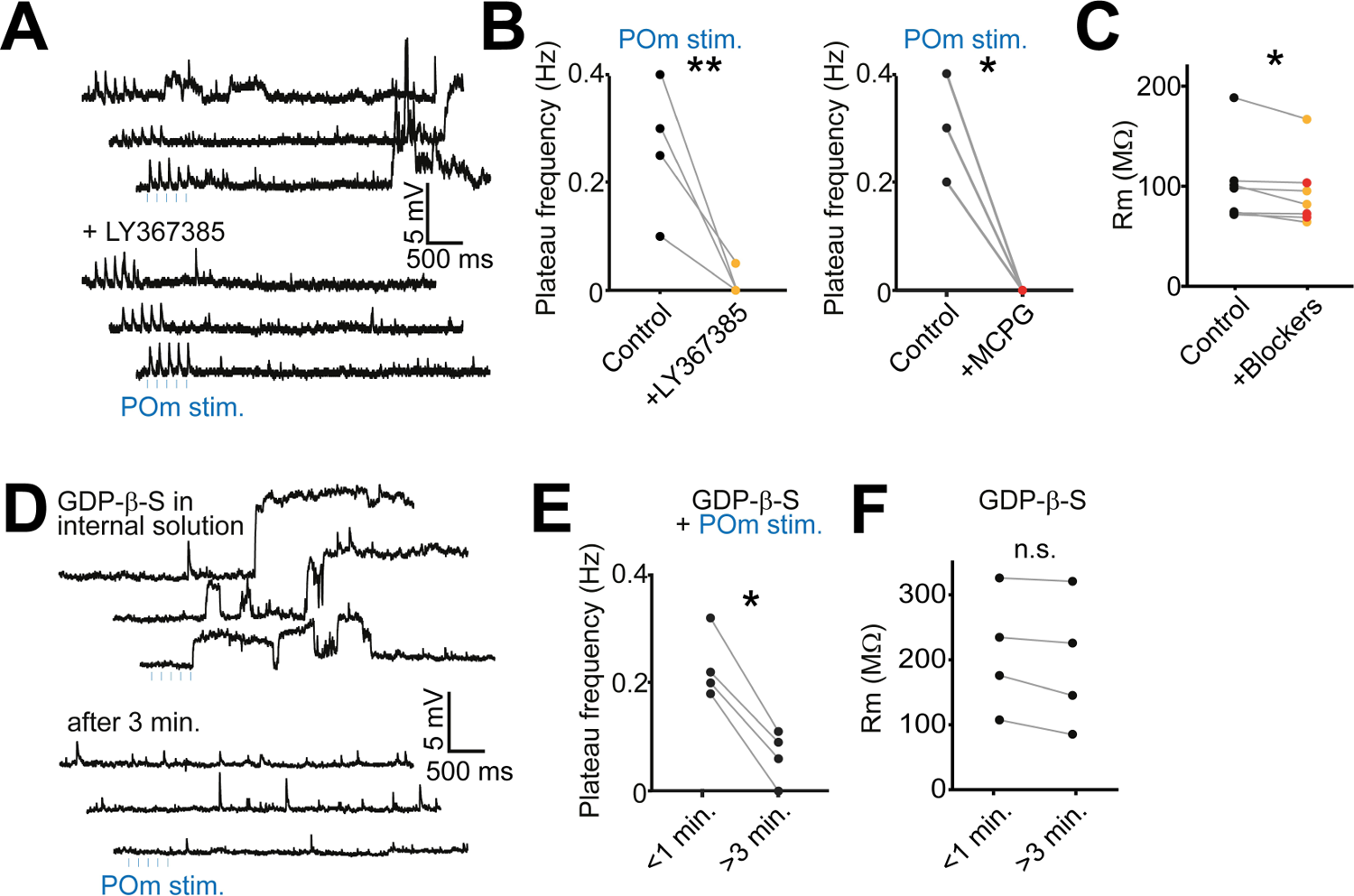
POm-mediated plateau potentials are mediated by the activation of mGluRIs (A) Dendritic recordings of a BT neuron during the stimulation of POm before and after the bath application of LY367385 (50 µM), a specific mGluRIs blocker. The drug prevented the generation of plateau potentials. (B) The plateau frequency was mostly prevented in the presence of LY367385 (50 µM; *n* = 4, *P* = 0.003, paired t-test) as well as in the presence of the generic mGluRIs blocker MCPG (500 µM; *n* = 3, *P* = 0.03, paired t-test). (C) Together, the mGluRI blockers LY367385 and MCPG significantly decrease the membrane resistance of the recorded dendrites (*n* = 7, *P* = 0.04, paired t-test). (D) Dendritic recording of a BT neuron during the stimulation of POm in the presence of GDP-β-S (1 mM), a G-protein activity blocker, in the intracellular solution. Within the first minute after break-in, plateau potentials could be observed but were not visible after 3 minutes, consistent with the dialysis of the drug. (E) The frequency of plateau potentials was largely reduced after intracellular dialysis of GDP-β-S (*n* = 4, *P* = 0.003, paired t-test). (F) The dialysis of GDP-β-S did not significantly reduce the membrane resistance of the dendrite (*n* = 4, *P* = 0.066, paired t-test).

### Modulation of mGluRIs alters movement-associated spiking of L2/3 neurons *in vivo*

mGluRI-mediated plateau potentials and the associated increase in input resistance may represent a mechanism for increasing the gain of concomitant synaptic inputs. Indeed, occasionally action potentials were superimposed on the plateau potentials (Figure 4A). To further investigate this, we performed dendritic recordings of BT neurons while bath applying TBOA, a glutamate-reuptake inhibitor that prolongs the presence of ambient glutamate in (and around) the synapse. Under these conditions, the optogenetic stimulation of POm afferents increased the occurrence of action potentials that were superimposed on the plateau potentials (Figure S10A,B). This effect was absent upon stimulation of M1 afferents, despite the amplifying effect of TBOA on the evoked PSPs (Figure S10C). This suggests that the plateau potentials are the leading cause for the increased occurrence of action potentials, which could trigger NMDAR-mediated events, as previously reported^52,53^. Affirmatively, the high amplitude plateau potentials and spikes disappeared when NMDARs were blocked by adding APV to the bath, but this did not impact the duration of the plateau potentials (Figure S10A,B). These could only be removed by an additional inhibition of mGluRI (Figure S10A,B). Altogether, the data indicate that under a prolonged presence of glutamate – a phenomenon that may mimic conditions as they occur during bursting activity, mGluRI-mediated plateau potentials may promote the generation of somatic action potentials.

These observations incited us to explore how the modulation of mGluRI affects the activity of cortical neurons *in vivo*. Mice actively use their whiskers to sense their environment, which consists of volitional movements that are in part initiated by activity in motor cortices, among which M1^54^. Neurons in the vibrissal area of M1 encode whisking parameters during active sensing behavior, and this activity is subsequently transmitted back to L1 of S1^55^. Based on our observations, we argued that POm-mediated activation of mGluRI could selectively increase the gain of incoming motor signals from M1 onto BT neurons. Therefore, we hypothesized that mGluRI-mediated subthreshold plateau potentials in S1 L2/3 pyramidal neurons increase the propensity for active whisking to induce somatic spikes. To investigate this, we performed *in vivo* 2-photon laser scanning microscopy to image calcium (Ca^2+^) signals in S1 L2/3 neurons expressing GCaMP6s. Ca^2+^ signals were recorded before and after modulating mGluRIs using the local infusion of the agonist DHPG or the antagonist MCPG (Figure 7). Using a piezo-driven microscope objective, we imaged near-simultaneously the upper and lower L2/3 neurons (at −100 and −300 µm distance from the pia, respectively; Figure 7A). For the analysis, we assumed that the population of BT neurons is enriched in upper L2/3 while the location of ST neurons is more biased towards deeper L2/3 (Figure 1E). We first compared the level of the overall activity of individual neurons in upper and lower L2/3 before and after DHPG (Figure 7B,D,E) or MCPG infusion (Figure 7C,F,G). DHPG increased the overall activity of L2/3 neurons but more so in upper L2/3 (Figure 7D,E). MCPG modestly increased the overall activity, but substantially suppressed activity of a subset of upper L2/3 neurons (Figure 7F,G). We tracked snout movements as a proxy of whisking^56^ using DeepLabCut^57^. A random forests decoding algorithm was trained to predict snout movements from the activity of individual neurons (Figure 7H,I). The correlation coefficient between the predicted and actual movements, which we defined as the prediction power (PP), was calculated for each neuron before and after infusion of the drugs (Figure 7J-M). While the average PP of upper and lower L2/3 neurons was similar in baseline conditions, we found that DHPG significantly increased the PP for upper but not for lower L2/3 (Figure 7J,K). Conversely, MCPG significantly decreased the PP of upper but not of lower L2/3 neurons (Figure 7L,M). In addition, the PP of upper L2/3 neurons under control conditions moderately correlated with the magnitude of increase in activity under DHPG, and strongly correlated with the reduction in activity under MCPG. This was not observed for lower L2/3 neurons (Figure S11). Altogether, these results indicate that mGluRI signaling in a subset of upper L2/3 neurons increases their propensity to produce spikes during whisking.

**Figure 7.**
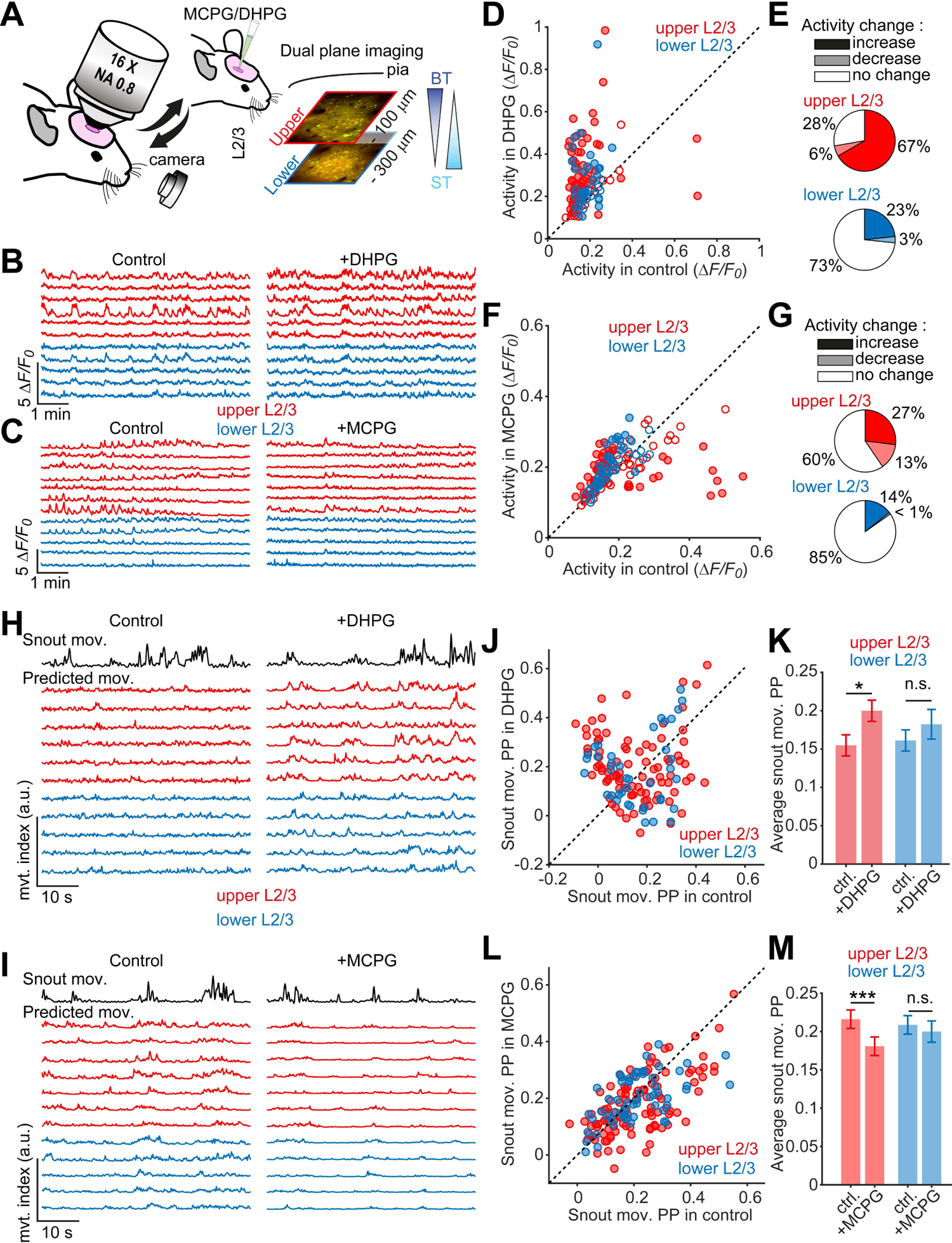
Modulation of mGluRIs *in vivo* bidirectionally changes the integration of movement-related information in upper L2/3 neurons (A) *In vivo* 2-photon calcium imaging was performed before and after injection of DHPG or MCPG in the barrel cortex directly through the silicone port of the cranial window. Upper and lower L2/3 neurons expressing GCaMP6s and mRuby2 were recorded quasi-simultaneously at −100 and −300 µm, respectively, from the pial surface. (B) Example traces of upper (red) and lower (blue) L2/3 neurons before and after injection of DHPG. (C) Same as (B), before and after injection of MCPG. (D) Comparison of the average normalized activity during the baseline recording and after DHPG injection for the upper (red, *n* = 90 neurons) and lower (blue) L2/3 neurons (*n* = 40 neurons, from 5 mice). Neurons that changed their activity level (closed circles) were determined with a permutation test (*P* < 0.01), shuffling data points between baseline and after drug injection and exhibiting at least a small effect size (Cohen’s *d* > 0.2). Other neurons were considered to not change their activity level (open circles). (E) Fractions of neurons showing an increase, decrease or no change in activity level after DHPG injection for the upper (red) and lower (blue) L2/3 neurons. (F) Same analysis as (D) for MCPG injection (*n* = 104 upper and 61 lower L2/3 neurons, from 3 mice). (G) Fractions of neurons showing activity change after MCPG injection. (H) and (I) A random forests model was used to evaluate the ability of individual L2/3 neurons to predict the snout movement before and after DHPG (H) or MCPG (I) injection. Predicted snout movements of the neurons shown in (B) and (C) were displayed (upper L2/3 neurons in red and lower L2/3 neurons in blue) and compared to the actual snout movement (top black trace). (J) Comparison of the snout movement PP (determined as the Pearson’s correlation between predicted and actual traces) in baseline and after DHPG injection (*n* = 90 upper and 40 lower L2/3 neurons, from 5 mice). (K) Average snout movement PP in baseline and after DHPG injection for the upper (red) and lower (blue) L2/3 neurons. Snout movement PP increased for the upper L2/3 neurons (paired t-test, *P* = 0.026) but not for the lower L2/3 neurons (*P* = 0.44). (L) Same analysis as (J) but for MCPG injection (*n* = 104 upper and 61 lower L2/3 neurons, from 3 mice). (M) Same as for (K) for MCPG injection. Snout movement PP decreased for the upper L2/3 neurons but not for the lower L2/3 neurons (for upper L2/3 neurons: *P* = 4e-4; for lower L2/3 neurons: *P* = 0.46; paired t-tests).

## DISCUSSION

We showed that activation of thalamocortical projections to S1 L2/3 pyramidal neurons from the POm promotes the generation of NMDA spikes when combined with other inputs. POm thalamocortical inputs also have the capacity to evoke plateau potentials via a mGluRI-mediated modulation of K2P channels. These effects are selective for a subpopulation of pyramidal neurons with broad apical tufts which are predominantly located in L2. Both phenomena actively increase neuronal excitability, which may augment these neurons’ propensity to trigger spikes. We found support for this mechanism *in vivo* by demonstrating that mGluRI signaling preferentially modulates movement-associated spiking of L2 pyramidal neurons in S1.

### POm thalamocortical projections preferentially connect with L2/3 BT neurons

Using an unbiased classification, we clustered L2/3 pyramidal neurons in two groups, one with broad and dense apical tufts, and another with slender tufts (Figure 1). The morphological characteristics of these populations are very similar to previously reported BT and ST neurons^8,9^. Our experiments show that the extent and density of their dendrites were good indicators of the synaptic connectivity with the axons that overlapped with them (Figure 2), which is in line with Peters’ rule stating that synaptic connections are proportional to the axo-dendritic overlap^12^. Stimulation of POm, M1 and S2 axons all tended to evoke larger PSPs in BT as compared to ST neurons, although the S2 and M1 functional inputs were not significantly different between both cell types. In contrast, stimulation of VPM and intracortical circuits evoked larger PSPs in the ST neurons which have a higher proportion of basal dendrites. We verified that this relationship was also present at the level of monosynaptic inputs between these neurons and POm or VPM axons (Figure S3). We did not perform this experiment for S2 and M1 afferents. Therefore, we cannot exclude that the lack of discrimination between those inputs onto BT and ST neurons was due to abundant polysynaptic connections from local corticocortical circuitry that may be activated by these pathways. The relatively strong connectivity between L1 afferents and BT neurons is in line with the notion that many L2 neurons in S1 and L2/3 neurons in V1 bearing complex dendritic arbors have relatively large receptive fields, since many L1 inputs derive from long-range projections originating from various brain regions^7,25,58^.

The high connectivity rates of POm afferents with L2/3 BT neurons stand in contrast with the relatively low connectivity rates of those afferents with the abundantly present branches from broad tufted L5b neuron apical dendrites in L1, and the high rates with less abundant slender L5a neurons^4^. Thus, POm afferents provide input selectively to L2 BT neurons and L5a pyramidal neurons, which are highly interconnected^1,3,16^. This suggests that together they constitute a paralemniscal cortical circuit motif with distinct functions^59^. It is also interesting to note that the high connectivity rates of L2 neurons (i.e. L2/3 BT neurons) with POm afferents are associated with higher-than-average levels of plasticity^25,59,60^. However, there is no indication that the BT neuronal subtypes should generally display higher levels of activity *in vivo*^9^.

### Converging POm and M1 inputs on L2/3 BT neurons cooperate to generate NMDA spikes

NMDA spikes were readily generated in BT neurons when POm afferents were co-stimulated with inputs from M1 and VPM, but not upon co-stimulation of VPM and S1_intracortical_ afferents (Figure 3). Conversely, NMDA spikes were abundant in ST neurons upon co-stimulation of VPM and S1_intracortical_ afferents but not when POm afferents were co-stimulated with those from M1 or VPM. Co-stimulation of POm and S2 afferents did not produce a significant number of NMDA spikes in either BT or ST neurons. The efficacy at which POm together with M1, and VPM together with S1_intracortical_, evoked NMDA spikes in BT and ST neurons, respectively, indicates that their synapses bolster supralinear synaptic interactions. Well known parameters for such interactions are synaptic proximity, the caliber of the parent dendrite, synaptic receptor content, the temporal order of activation, and the levels of local inhibitory and neuromodulatory input^37^. Favoring conditions are indeed met for the synaptic inputs of some of the above afferents. POm and M1 afferents both are likely to have dense connections with the apical dendrites of BT pyramidal neurons in L1, and VPM and S1_intracortical_ afferents may densely connect with basal dendrites of ST pyramidal neurons in L3. Thus, the occurrence of NMDA spikes may be related to the convergence of these inputs onto single dendritic domains, which aligns with models in which clustered inputs favor supralinear synaptic integration^61–65^. In this respect, the finding that co-stimulation of POm and VPM afferents also evoked NMDA spikes in BT neurons was surprising, since VPM axons had virtually no monosynaptic connections with them. The most likely explanation for this finding is that they were triggered by the activation of polysynaptic L4-to-L2/3 and L2/3-to-L2/3 circuits that are readily recruited by VPM inputs, and which are more broadly distributed along the dendrites. In addition to their convergence, another favoring condition for NMDA spikes is provided by the location of POm synaptic inputs on thin distal endings of the apical branches^66^, which may increase their cooperativity with other inputs^42^. In support of this, NMDA-mediated synaptic Ca^2+^ responses can readily be observed in distal dendritic branches ^23,39^. In a similar vein, the generation of NMDA spikes in ST neurons could be explained by the projection of VPM afferents onto their thin basal dendrites which are also favorable for evoking dendritic spikes^67,68^. A similar supra-linear interaction between VPM and local inputs has been described for L4 granule cells^38^. Furthermore, co-stimulation of POm afferents with local ascending cortical circuitry has been shown to drive disinhibition of L2/3 neurons^26^. This could aid the generation of NMDA spikes, which are highly sensitive to dendritic inhibition^69^. Modeling studies indeed suggest that disinhibition could gate the generation of NMDA spikes evoked by clustered inputs^70^. L2 pyramidal neurons receive substantial inhibition and could thus be powerfully controlled by such a mechanism^71^. Lastly, the various inputs that we tested could harbor temporally different synaptic activation patterns, which have been shown to be critical for supra-linear synaptic integration^72^. Further experiments are needed to reveal if such relationships exist between long-range synaptic inputs to apical dendrites in L1.

Even though NMDA spikes often remain below the main spiking threshold, they are important for facilitating spikes initiated at the axon initial segment. Dendritic NMDAR-mediated events *in vivo* have been correlated with increased somatic spiking probabilities^38,39,73^. Thus, POm afferents may modulate activity of BT neurons through facilitating the generation of NMDA spikes in collaboration with other inputs. Subthreshold NMDAR-events have also been shown to drive synaptic plasticity^23,43,74^, and previous work from our laboratory indicates that in L2/3 neurons this depends on input from the POm. Combined with the current insights this suggests that POm-mediated plasticity might be an attribute of L2/3 BT neurons. It will be interesting to investigate whether this relates to aspects of sensory learning.

### Stimulation of POm afferents modulates K2P leak channels through mGluRI signaling

POm stimulation *in vivo* has been shown to evoke sustained depolarizations in L2/3 and L5 neurons *in vivo*, which in turn may prolong their sensory-evoked responses^23–25,27^. Reverberatory activity in L2 is likely evoked through a combination of L5-to-L2 and direct POm-to-L2 inputs^25,59^. POm-mediated NMDA spikes may in part explain such effects, but additional biophysical mechanisms for these phenomena are likely at play given the long-lasting effects seen in the studies above (Figure S10). Upon a burst of POm stimuli we observed abrupt depolarizations specifically in BT neurons, which were temporally unlocked from the stimulus onset and varied in duration (Figure 4). These potentials were very distinct from PSPs and synaptic NMDA spikes (Figure S7). Instead, they had similar kinetics to previously reported local glutamate-evoked dendritic plateau potentials that were sustained at 10 to 20 mV for 200-500 ms^53,75^. In these studies, the potentials propagated to the cell body where they could trigger a burst of action potentials, an effect that we did not further investigate in brain slices. Surprisingly, in our experiments, we found that plateau potentials were mediated by mGluRI, which modulated K2P channels through a G-protein-triggered mechanism (Figure 5 and 6). Our results point to a mechanism involving TASK and TREK K2P channels, but other channels could play a role as well since our pharmacological screen could not exhaustively test specific interactions.

Although this mechanism may seem surprising, it does align with other observations. First, synaptic responses elicited by strong higher-order thalamocortical stimuli were shown to consistently contain mGluRI components^21,22^. Second, activation of mGluRs in neocortex and hippocampus is known to modulate various K^+^ channels^76–81^, although Ca^2+^ and other channels can also be affected^82^. Third, signaling pathways from mGluRs to K2P channels have been identified using reduced expression systems^83,84^ and in cerebellum granule cells^47,48,85^.

K2P channels are responsible for leak and background currents, which are largely voltage-independent^86,87^. As such, they play an important role in regulating the resting membrane potential and hence a neuron’s excitability. By reducing the number of background potassium channels that are in an open state, mGluR signaling could depolarize the membrane by a few mV. The metabotropic action of the receptors together with their extrasynaptic location might make this a delayed, slow, yet all-or-nothing effect. The delays and the kinetics of the plateau potentials that we detected upon POm stimulation are consistent with such a mechanism. It is important to note that the interaction between mGluR and K2P are intricate, and may involve antagonistic signaling cascades^47,48^. It will be interesting to investigate if the various interactions depend on the molecular composition and if they are selective for synaptic or neuronal types. Furthermore, other GPCR-signaling pathways may also converge onto the K2P channels, e.g. through cholinergic signaling^48^.

Our observation that the mGluRI-mediated modulation of the K2P channels only occurred for POm-to-BT inputs, and not by neighboring long-range synaptic inputs, suggests that selectivity is regulated at the synaptic level. A combination of postsynaptic scaffold proteins may selectively recruit mGluRs, or their precise localization could be regulated through trans-synaptic interactions similar to the recruitment of presynaptic mGluR7^88^ – although such interactions have as yet not been described for mGluR1 or 5. Synapse selectivity could also emerge from differential expression and trafficking of K2P family members or the intermediate signaling molecules^87,89^ – although it is not clear whether localization can also be dendritic domain-specific.

### The role of POm-mediated mGluR signaling in regulating excitability

The POm evoked mechanisms could increase the excitability of pyramidal neurons in two ways. First, the dendritic plateau potentials bring the resting membrane potential into a more depolarized state, which may increase the probability that coinciding EPSPs cross the spiking threshold. In this respect, the POm-driven plateau potentials are comparable to the cortical up-states that were previously reported upon optogenetic stimulation of thalamocortical circuits^90^. Second, the modulation of K^+^ channels may facilitate the transformation of electrogenic events originating in individual dendrites into global dendritic activity (i.e. simultaneous depolarizations in many dendrites)^33,91^, which may promote the generation of action potentials^92–95^. Indeed, dendritic K^+^ conductance has been shown to inhibit the initiation of local supralinear events, prevent the backpropagation of action potentials into the dendrites, dampen excitatory synaptic events, and more generally, decouple the dendritic from the somatic compartment^94,96^. Interestingly, thalamic activation (POm) of mGluRI has been found to be necessary for the coupling between the dendrites and soma of L5 pyramidal neurons^97,98^. Together with our findings, these observations suggest that a POm-driven block of K2P-conductance increases the excitability of distal dendritic compartments and thereby increases the transfer of depolarizations caused by other distal inputs from the dendrites to the soma. Overall, this aligns with studies showing that the activation of the POm causes a general increase in cortical excitability of the barrel cortex^24,66^, and that the activation of postsynaptic mGluR5 receptors induces persistent firing in the prefrontal cortex^99^. Also, in line with these findings, we found that the local injection of mGluRI agonists and antagonists *in vivo* bidirectionally affected movement-associated activity of L2 neurons. In particular, the movement-related prediction from L2 neurons’ activity increased upon the presence of mGluRI agonists and decreased with antagonists (Figure 7). This strongly suggests that L2 pyramidal neuron activity, which likely depends on active feedback loops among others from POm, is mediated by mGluRI-associated mechanisms; possibly through the modulation of K2P channel opening. The regulation of excitability and dendritic coupling of various pyramidal neurons might rely on this mechanism. Recent work by the Larkum laboratory has shown that anesthetics cause dendritic decoupling and propose that this could be the underlying mechanism for the loss of consciousness^98^. Interestingly, K2P channels have been found to be positively modulated by some anesthetics^100,101^. This implies that POm inputs to L5 and L2/3 may play an important role in modulating levels of consciousness, which could even be a general feature of higher-order thalamocortical inputs to various cortical areas.

## ACKNOWLEDGMENTS

We would like to express our gratitude to Dr. Fodoulian for generously providing the random forests code, which significantly enriched the computational aspects of our manuscript. We also want to extend the appreciation to the entire Holtmaat group for their valuable comments and constructive feedback, which have been instrumental in shaping and refining the development of this project. We would like to thank Elodie Husi, Sébastien Pellat and Raphaël Thurnherr for their technical support. This work was supported by the Swiss National Foundation Grant Numbers 31003A_173125 and 310030_204562.

## AUTHOR CONTRIBUTIONS

Conceptualization, F.B., R.C. and A.H.; Methodology, F.B., R.C., F.M., S.P. and A.H.; Investigation, F.B., R.C., C.M. and N.M.; Formal Analysis, F.B., R.C., T.B.; Resources, A.H.; Writing – Original Draft, F.B., R.C. and A.H.; Writing – Review & Editing, F.B., R.C. and A.H.; Visualization, F.B., R.C. and A.H.; Funding Acquisition, A.H.; Supervision, A.H.

## STAR METHODS

### KEY RESOURCES TABLE

| REAGENT or RESOURCE | SOURCE | IDENTIFIER |
| --- | --- | --- |
| Viral vectors |  |  |
| AAV2/5-EF1a.eGFP.WPRE.RBG | University of Pennsylvania Vector Core | Cat # 105547-AAV5<br>RRID: Addgene_105547 |
| AAV2-CB7.Cl.mCherry.WPRE.RBG | University of Pennsylvania Vector Core | Cat # 105544-AAV2<br>RRID: Addgene_105544 |
| AAVrg-pmSyn1.EBFP-Cre | AAV pmSyn1-EBFP-Cre was a gift from Hongkui Zeng (http://n2t.net/addgene:51507) | Addgene(51507-AAVrg),<br>RRID: Addgene_51507 |
| pAAVrg-CAG-tdTomato | pAAV-CAG-tdTomato (codon diversified) was a gift from Edward Boyden (http://n2t.net/addgene:59462) | Addgene(59462-AAVrg),<br>RRID: Addgene_59462 |
| AAV1-Flex-hSyn1-mRuby2-GSG-P2A-GCaMP6s-WPRE-pA | pAAV-hSyn1-Flex-mRuby2-GSG-P2A-GCaMP6s-WPRE-pA was a gift from Tobias Bonhoeffer & Mark Huebener & Tobias Rose (http://n2t.net/addgene:68720) | Addgene(68720-AAV1),<br>RRID: Addgene_68720 |
| AAV9-CaMKII-0.4.Cre-SV40 | pENN.AAV.CamKII 0.4.Cre.SV40 was a gift from James M. Wilson (http://n2t.net/addgene:105558) | Addgene(105558-AAV9),<br>RRID: Addgene_105558 |
| AAV1-mCaMKIIα-iCre-WPRE-hGHp(A) | UZH Viral Vector Facility | v206-1 |
| AAV2-hSyn-DIO-eGFP | pAAV-hSyn-DIO-EGFP was a gift from Bryan Roth (http://n2t.net/addgene:50457) | Addgene(50457-AAV2),<br>RRID: Addgene_50457 |
| AAV2-CaMKIIα-hChR2(H134R)-eYFP | pAAV-CaMKIIα-hChR2(H134R)-EYFP was a gift from Karl Deisseroth (http://n2t.net/addgene:26969) | Addgene(26969-AAV2),<br>RRID: Addgene_26969 |
| AAV2-EF1a-DIO-hChR2(E123T/T159C)-eYFP | pAAV-Ef1a-DIO hChR2(E123T/T159C)-EYFP was a gift from Karl Deisseroth (http://n2t.net/addgene:35509) | Addgene(35509-AAV2),<br>RRID: Addgene_35509 |
| Chemicals, peptides, and recombinant proteins |  |  |
| (RS)-3,5-DHPG | Tocris Bioscience | Cat. No. 0342 |
| (RS)-MCPG | Tocris Bioscience | Cat. No. 0336 |
| APV | Sigma Aldrich | A8054 |
| Bupivacaine hydrochloride | Sigma Aldrich | PHR1128 |
| QX 314 chloride | Tocris Bioscience | Cat. No. 2313 |
| Barium chloride dihydrate | Sigma Aldrich | 217565 |
| A1899 | Tocris Bioscience | Cat. No. 6972 |
| Fluoxetine hydrochloride | Tocris Bioscience | Cat. No. 0927 |
| IBMX | Tocris Bioscience | Cat. No. 2845 |
| LY367385 | Tocris Bioscience | Cat. No. 1237 |
| TETRODOTOXIN Citrate (TTX) | Latoxan | L8502 |
| 4-Aminopyridine (4AP) | Sigma Aldrich | A78403 |
| NBQX disodium salt (NBQX) | Abcam | ab120046 |
| (+)-Bicuculline | Tocris Bioscience | Cat. No. 0130 |
| DL-TBOA | Tocris Bioscience | Cat. No. 1223 |

| Deposited data |  |  |
| --- | --- | --- |
| Raw and analyzed data | This study | https:// doi.org/ 10.5281/zenodo.10210325 |
| Experimental models: Organisms/strains |  |  |
| Mouse/C57BL/6JRj | Janvier Labs | https://janvier-labs.com/fiche_produit/2-c57bl-6jrj/ |
| Mouse/ Pvalbtm1(cre)Arbr/J (PVcre) | The Jackson Laboratory | MGI:3590684<br>https://www.jax.org/strain/008069, RRID: IMSR_JAX:008069 |
| Software and algorithms |  |  |
| Fiji | Schindelin et al.<sup>102</sup> | https://imagej.net/Fiji |
| pClamp 10.5 | Molecular Devices, LLC | https://www.moleculardevices.com/ |
| Clampfit 10 | Molecular Devices, LLC | https://www.moleculardevices.com/ |
| Origin 2021 | OriginLab Corporation | https://www.originlab.com/2021 |
| Prism | GraphPad | https://www.graphpad.com/ |
| Neurolucida | MBF Bioscience | https://www.mbfbioscience.com/products/neurolucida/ |
| Scanimage2016 | Vidrio Technologies<br>Pologruto et al.<sup>103</sup> | https://www.mbfbioscience.com/products/scanimage/ |
| Matlab2018a and 2020b | The MathWorks, Inc | https://www.mathworks.com/products/matlab.html |
| Python | Python Software Foundation | http://www.python.org/ |
| DeepLabCut v2.2.0.2 | Nath et al.<sup>104</sup> | https://github.com/DeepLabCut/DeepLabCut |
| R v4.0.3 | The R Foundation | http://www.r-project.org/ |

### RESOURCE AVAILABILITY

#### Lead contact

Further information and requests for resources and reagents should be directed to and will be fulfilled by the lead contact, Anthony Holtmaat.

#### Materials availability

This study did not generate new unique reagents.

#### Data and code availability

- The data used to generate the figures is freely available at the CERN data repository Zenodo https://zenodo.org/communities/holtmaat-lab-data/ with https://doi.org/10.5281/zenodo.10210325.
- The principal Matlab code that was used for data analysis is freely available at the CERN data repository Zenodo https://zenodo.org/communities/holtmaat-lab-data/ with https://doi.org/10.5281/zenodo.10210325.

## EXPERIMENTAL MODEL AND SUBJECT DETAILS

### Mice

Male C57BL/6J wild-type (Charles River or Janvier Labs) and PV-Cre mice (https://www.jax.org/strain/017320), aged 8 to 12 weeks, were group housed with littermates on a normal 12-h light cycle with food and water available *ad libitum*. All procedures were conducted in accordance with the guidelines of the Federal Food Safety and Veterinary Office of Switzerland and in agreement with the veterinary office of the Canton of Geneva (license numbers GE12219B, GE/74/18 and GE253A).

## METHOD DETAILS

### Virus injection for electrophysiology

C57BL/6J or Parvalbumin (PV)-Cre mice, 8–12 weeks old, were anesthetized with isoflurane mixed with oxygen (3–5% induction, 1–2% maintenance), placed in a stereotaxic apparatus, and prepared for injections with craniotomies over the target injection regions. Deep anesthesia was assessed by absence of foot pinch reaction. The skin overlying the skull was removed under local anesthesia using Carbostesin (AstraZeneca) or Lidocaine (Streuli). Mice were then head-fixed with ear-bars and a nose clamp on a stereotaxic apparatus (Stoelting). Eyes were protected from drying with artificial tears. The body temperature was monitored with a rectal probe and was maintained at ∼37°C using a heating pad (FHC) during surgery. Bilateral craniotomies were performed using an air-pressurized driller and injections (100-200 nl per injection site) were performed using a pulled glass pipette (10–15 µm diameter tip) mounted on a Nanoject II small-volume injector (Drummond Scientific). Injections were performed at a speed of 23 nl/s, separated by 2-3 min intervals, in POm (2.2 mm posterior to bregma, 1.2 mm lateral and 3 mm below the bregma), VPM (1.85 mm posterior to bregma, 1.75 mm lateral and 3.5 mm below the bregma), S1 (1.5 mm posterior to bregma, 3.5 mm lateral and 0.4 mm below the pial surface), M1 (1.54 mm anterior to bregma, 1.75 mm lateral and 0.5 mm below the pial surface) and S2 (0.7 mm posterior to bregma, 4.2 mm lateral and 0.3 mm below the pial surface). Different viruses (AAV2/5-EF1a.eGFP.WPRE.RBG; AAV2-CB7.CI.mCherry.WPRE.RGB; AAVrg-pmSyn1.EBFP-Cre; pAAVrg-CAG-tdTomato; AAV2-rAAV.EF1a-DIO-hChR2(E123/T159C)-eYFP; AAV2-CaMKIIα-hChR2-eYFP) were injected with regards to the different experiments. All injections were bilateral. The pipette was left in place for 3–5 min before removing it from the brain. Mice were given analgesics (carprofen 5 mg/kg; TW Medical, #PF-8507) after surgery and monitored daily to ensure full recovery. Animals were then put back in their home cage to recover from the surgery. A minimum period of three weeks was allowed for viral expression before the animals underwent additional experimental procedures.

### Acute brain slice preparation

Mice were anesthetized with a ketamine/xylazine (100 mg/kg, 10 mg/kg) cocktail and were perfused intracardially with ice-cold high sucrose saline solution consisting of the following (in mM): 2.8 KCl, 1.25 NaH_2_PO_4_, 25 NaHCO_3_, 0.5 CaCl_2_, 7 MgCl_2_, 7 dextrose, 205 sucrose, 1.3 ascorbate, and 3 sodium pyruvate (bubbled with 95% O_2_/5% CO_2_ to maintain pH at ∼7.4). A vibrating tissue slicer (Leica VT S1000, Germany) was used to make 250-μm-thick sections from 0.58 to 1.46 mm posterior to the bregma position. For obtaining S1 acute slice, the brain was removed and mounted to the stage of the vibratome, and sections were made coronally. Slices were held for 30 minutes at 35°C in a chamber filled with aCSF consisting of the following (in mM): 125 NaCl, 2.5 KCl, 1.25 NaH_2_PO_4_, 25 NaHCO_3_, 2 CaCl_2_, 2 MgCl_2_, 10 dextrose, and 3 sodium pyruvate (bubbled with 95%O_2_/5% CO_2_) and then at room temperature until the time of recording.

### Whole cell recording

The intracellular solution contained the following (in mM): 120 K-gluconate, 16 KCl, 10 HEPES, 8 NaCl, 7 Mg^2+^-phosphocreatine, 0.3 Na-GTP, 4 Mg-ATP, pH 7.3 with KOH^105,106^. Biocytin (Vector Laboratories; 0.1%-0.2%) was also included for histological processing and *post hoc* cell location determination. In some experiments, Alexa-594 (16 μM; Thermo Fisher Scientific, #A10428) was also included in the internal recording solution to determine the dendritic recording location relative to the soma as well as a first assessment of the cell morphology. Dendritic recordings were performed at approximately 157 ± 25 µm from the soma for BT neurons and 208 ± 43 µm for ST neurons. Data was acquired using a Multiclamp 700b amplifier and the Clampex11 (Molecular Devices) data acquisition software. Data were acquired at 10–50 kHz, filtered at 2–10 kHz, and digitized by an Axon Digidata 1550B interface (Molecular Devices). Pipette capacitance was automatically compensated for. Series resistance was monitored and compensated throughout each experiment and was 10-25 MΩ for somatic recordings and 15–40 MΩ for dendritic recordings. Recordings were discarded if series resistance increased by more than 30% during the recording. Voltages are not corrected for the liquid-junction potential (estimated as ∼8 mV). The acute slice was placed in a recorded chamber with a feedback temperature system set at 36 degrees and continuously perfused with oxygenated ACSF (flow rate 1.5 ml/min). For optogenetic stimulation, 5-ms long light pulses were delivered through the objective using the coolLED pE-300ultra (CoolLED Ltd.), delivering blue light (475 ± 23 nm) for activating ChR2 and/or amber light for activating ChrimsonR (575 ± 25 nm). The intensity of the LED was normally set to 7 mW/mm^2^ (10% of the maximum LED power) for the blue light and 6.3 mW/mm^2^ (30% of the maximum LED power) for the amber light unless stated otherwise in the text.

Depending on the experiments, the following drugs were perfused in the bath, together or sequentially as described in the main text and figure legends: APV (50 µM, Sigma Aldrich), bupivacaine (1 mM, Sigma Aldrich), QX-314 (1 mM, Tocris Bioscience), Barium (1 mM, Sigma Aldrich), A1899 (100 nM, Tocris Bioscience), Fluoxetine hydrochloride (100 µM, Tocris Bioscience), IBMX (200 µM, Tocris Bioscience), LY367385 (50 µM, Tocris Bioscience), MCPG (500 µm, Tocris Bioscience), TTX (1 µM; Latoxan), 4AP (100 µM; Sigma Aldrich), NBQX (10 µM, Abcam), bicuculline (10 µM, Tocris Bioscience), DHPG (10 µM, Tocris Bioscience), TBOA (10 µM, Tocris Bioscience). For some experiments, GDP-β-S (1 mM, Sigma Aldrich) was mixed with the intracellular solution.

### Analysis on electrophysiological recordings

The parameters of the optogenetically evoked events, such as the rise time, amplitude, duration, were analyzed using built-in functions in Clampex (Molecular Devices).

To assess if a neuron exhibited dSpikes with the co-stimulation of a pair of inputs, we analyzed the distribution of the mean amplitude of the optogenetically evoked events and calculated a bimodality coefficient from the distribution as follow:

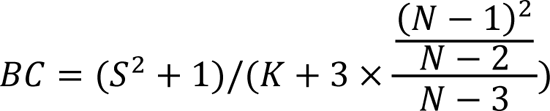

where *N* is the number of samples, *K* and *S* are the data kurtosis and skewness respectively, calculated using MATLAB functions (Mathworks). We considered that a distribution was bimodal when *BC* was greater than 0.5. Subsequently, a *k*-means cluster analysis with *k* = 2 was applied to determine which trials displayed dSpikes, and this served to calculate the fraction of trials and the mean amplitude of dSpikes. In addition, dSpike strength was calculated by multiplying the fraction of trials with dSpikes with their mean amplitude.

Post-stimulation events were automatically identified by detecting the changes in the membrane potential using a threshold that corresponded to 3 times the standard deviation of the baseline noise. This baseline noise was defined as the data below the median value of the whole trace. Traces were then low pass filtered at 100 ms and events were detected as above the threshold for at least 10 ms. The amplitude and duration of all the detected events were then extracted. We used a conservative threshold of 200 ms to separate short and long-lasting events; the latter being classified as plateau potentials confirmed by visual inspection.

### Morphological identification and reconstruction of recorded neurons

When approaching with the path pipette, L2/3 pyramidal neurons were discriminated by the orientation of their main apical dendrite visualized in bright field mode. A cell with an oblique apical dendrite would be selected as a putative BT neuron. On the contrary, a cell with a main apical dendrite going straight toward the pia would be classified as a putative ST neuron.

Once the electrophysiological recording was completed, the electrode was gently pulled back from the dendrite to avoid membrane ruptures to let the recorded neuron retrospectively reconstructed using Neurolucida to confirm their dendritic morphology (Figure 1A, Figure S1A). More in detail, each slice was then transferred in paraformaldehyde (PFA) 4% in 0.1 M phosphate buffer saline (PBS) for 10-15 min, then stored in 0.1 M PBS at 4 °C until the beginning of the biocytin staining procedure (up to 1 week). As previously reported^107^, slices were washed in PBS, then incubated in 1% Triton for 30 min and in 0.5% H_2_O_2_ for 30 min. Slices were washed again in PBS and incubated with VECTASTAIN Elite ABC Horseradish Peroxidase kit (Vector Laboratories) for 48 hours at 4° C. Slices were washed again in PBS and reacted with the chromogen 3,3’-diaminobenzidine (DAB kit, Vector Laboratories). When the reaction was complete, slices were mounted with the Vectashield mounting medium (Vector Laboratories). Dendritic arborizations were reconstructed in bright field under a 100/1.30 NA oil-immersion objective using a Neurolucida system (MicroBrightField). Only spiny neurons were included in the reconstructed pool. VIP spiny neurons^108^ were discriminated based on their distinct electrophysiological passive and firing properties. Quantification of the length and number of branches was automatically extracted from the reconstructions. Basal and apical dendrites were defined automatically as all the dendrites that originate from below or above the centroid of the soma respectively (Figure S1).

### Surgery for in vivo calcium imaging

Stereotaxic injections of adeno-associated viral (AAV) vectors were carried out on 6 weeks old male C57BL/6J mice (Janvier Labs). Anesthesia was first induced by a mix of O_2_ and 4% isoflurane at 0.4 L.min^-1^ followed by an intraperitoneal injection of MMF solution, consisting of 0.2 mg.kg^-1^ medetomidine (Dormitor, Orion Pharma), 5 mg.kg^-1^ midazolam (Dormicum, Roche), and 0.05 mg.kg^-1^ fentanyl (Fentanyl, Sinetica) diluted in sterile 0.9% NaCl. A mix of AAV1-Flex-hSyn1-mRuby2-GSG-P2A-GCaMP6s-WPRE-pA and AAV9-CaMKII-0.4.Cre-SV40 (Addgene, 68720-AAV1 and 105558-AAV9 respectively) with a ratio of 20:1 was delivered to L2/3 of the right S1 at the approximate location of the C2 barrel-related column (1.4 mm posterior to the bregma, 3.5 mm to the right, −0.3 mm below the pial surface). A 3-mm diameter cranial window, prepared with a silicone port^109^, was implanted, as described previously^110^. Imaging was performed after at least 2 weeks of viral expression.

### In vivo drug injections

After recording the spontaneous activity of L2/3 neurons in baseline conditions, the mGluRI agonist DHPG (50 mM^111,112^; Tocris Bioscience) or antagonist MCPG (500 µM^113^; Tocris Bioscience) was injected through the silicone port of the cranial window using a glass pipette. A volume of 100 nl was slowly injected right below the pia. Mice were left to recover for 15 min before placing them back under the microscope for recording.

### Two-photon laser scanning microscopy

We used a custom built 2-photon laser scanning microscope mounted onto a modular *in vivo* multiphoton microscopy system (https://www.janelia.org/open-science/mimms-10-2016) equipped with an 8-kHz resonant scanner and a 16× 0.8NA objective (Nikon, CFI75), and controlled with Scanimage 2016b^103^ (http://www.scanimage.org). Fluorophores were excited using a Ti:Sapphire laser (Chameleon Ultra, Coherent) tuned to λ = 980 nm at an approximate power of 25 mW. Fluorescent signals were collected with GaAsP photomultiplier tubes (10770PB-40, Hamamatsu) separating mRuby2 and GCaMP6s signals with a dichroic mirror (565dcxr, Chroma) and emission filters (ET620/60m and ET525/50m, respectively, Chroma). Prior imaging, mice were handled and accustomed to being head restrained under the microscope for 10-15 min over 4-5 days. Two imaging depths were acquired quasi-simultaneously at approximately 10 Hz using a piezo z-scanner (P-725 PIFOC, Physik Instrumente) for moving the objective over the z-axis. The two planes were set with a size of 350 × 350 µm (512 × 256 pixels) and positioned at 100 and 300 µm below the pia (i.e. the upper and lower L2/3).

### Image processing

Images were processed using custom-written MATLAB scripts and ImageJ (http://rsbweb.nih.gov/ij/). Lateral motion corrections were performed using the reference mRuby2 signal, from the red channel. Rigid lateral movement vectors were calculated using the NoRMCorre MATLAB toolbox^114^. Residual bidirectional scanning artifact vectors were calculated using a highest-pixel-line signal correlation between the two scanning directions on the entire frame. All calculated lateral motion corrections were applied on both the mRuby2 and GCaMP6s channels. For an unbiased extraction of the GCaMP6s fluorescence signals from individual neurons, regions of interest (ROIs) were drawn manually for each session based on neuronal shape using the mRuby2 signal. The fluorescence time-course of each neuron and channels were measured as the average of all pixel values within the ROI. Local neuropil signal was measured for each ROI and channels as the average of pixel values within an automatically defined ring of 15 µm width, 2 µm away from the ROI and excluding overlapping regions with surrounding ROIs. Residual axial movement corrections were applied using the fluctuations in the mRuby2 signal of the measured ROIs. To perform this correction, signal traces were initially filtered using an exponential moving average filter with a window size of 500 ms. Then, the mRuby2 signal trace (*FR*_cell measured_) was rescaled to the GCaMP6s (*FG*_cell measured_) signal trace by normalizing the values using their 8^th^ percentiles (*minR* and *minG* respectively) and their median values (*medR* and *medG* respectively) over a rolling window of 180 s as:

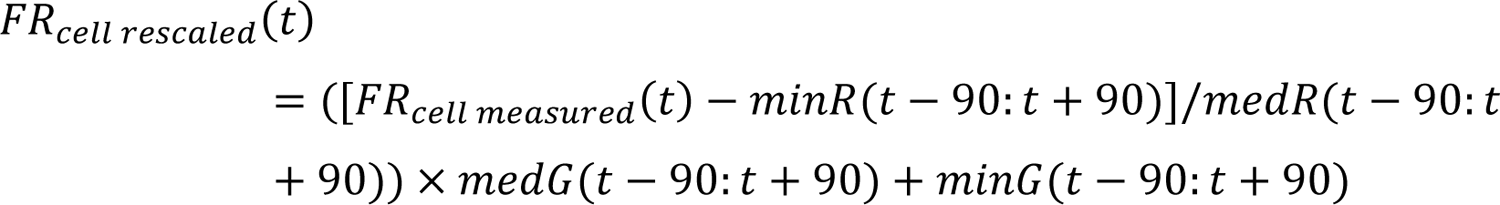

We used the median value for normalization to consistently compare the mRuby2 signal with basal GCaMP6s signal. The GCaMP6s signal was then corrected as follow:

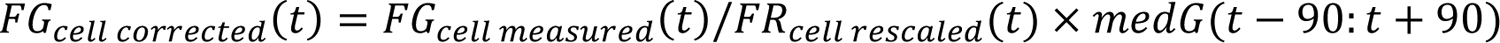

The same operations were performed on the neuropil signal to obtain the neuropil corrected vector (*FG*_neuropil corrected_). The true GCaMP6s signal of a cell body was then estimated as:

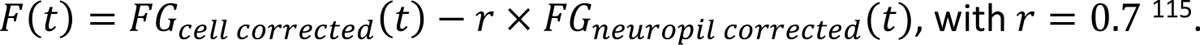

Normalized calcium traces Δ*F*/*F*_O_ were calculated as: (*F*(*t*) − *F*_O_)/*F*_O_, where *F*_O_ is the 30^th^ percentile of the whole *F* trace. To determine if a neuron had changed its level of activity after drug injection, we calculated the effect size (or Cohen’s d). The effect size corresponded to the difference in means between the before and after normalized calcium traces divided by the pooled standard deviation. To calculate the Cohen’s d, we used a permutation test by shuffling 1,000 times the datapoints between the two traces to create a null distribution of differences in means and standard deviations that would be expected under the null hypothesis of no difference between the two traces. We considered that a neuron changed its level of activity if the Cohen’s d was more than 0.2, which corresponds to a small effect size.

### Anterograde labeling and analysis

AAV injections were performed following a similar procedure as for surgeries for *in vivo* imaging but without cranial window implantation. For these experiments, 20 nl of AAV1-mCaMKIIα-iCre-WPRE-hGHp(A) was injected in either the VPM (1.8 mm posterior to bregma, 1.7 mm lateral and 3.5 mm below the bregma) or in the POm (2.2 mm posterior to bregma, 1.25 mm lateral and 3 mm below the bregma). A second injection of 200 nl of AAV2-hSyn-DIO-eGFP at the approximate C2 barrel coordinates (1.4 mm posterior to bregma, 3.5 mm to the right and 0.3 mm below the pial surface). After 3-5 weeks of viral expression, mice were perfused, and brain slices were cut at a thickness of 300 µm. Images stacks of 650 × 650 × ∼300 µm, with a voxel size of 0.3 × 0.3 × 1.5 µm, were acquired using 2-photon laser scanning microscopy (see above) tuned to 910 nm and a 25× 1.1NA objective (Nikon) at an approximate power of 50 mW. The positions of the soma were manually marked, and the pia position was automatically defined using a custom-written script in MATLAB.

### Decoding analysis

Snout movement recording was performed under 930 nm infrared illumination (M940L3, Thorlabs) using a 20 Hz infrared sensitive camera and the FlyCap acquisition software (FLIR Systems). Snout position was extracted from the raw video frames using the DeepLabCut tracking algorithm^57^. In brief, a model was created using hand-annotated sample frames of the position of the snout in different imaging sessions. The model was then applied to determine the position of the snout in each frame of all the videos. The movement was calculated as the sum of the absolute derivatives of the x and y positions coordinates and applied a low-pass filter at 1 Hz.

A random forests machine-learning algorithm was used to decode the snout movements from the activity of single neurons. Given the slow kinetics of calcium transients captured by the GCaMP6s sensor, spiking rates were inferred from the Δ*F*/*F*_O_ trace and used as input to the algorithm, which allowed to temporally match fast motor movements to neuronal activity. For this we used a fast nonnegative deconvolution method (https://github.com/jovo/oopsi)^116^ with variable background fluorescence estimation and a K_D_ of 144 nM^117^. For the algorithm to capture differences in activity levels between neurons, the activity traces from the movies before and after drug injections and of all neurons recorded were concatenated before inferring spikes. Both neuronal activity and behavior traces were resampled at 20 Hz. To account for putatively preceding pre-motor and/or following sensory-related activity in S1 relative to behavioral events, the neuronal activity traces were shifted negatively and positively in time with a maximum shift of 250 ms. Thus, eleven time-shifted inferred firing rate traces (discretized in time bins of 50 ms) centered on zero time shift were used to predict instantaneous behavioral features and composed a vector *X*_i_(*t*) = [*x*_i_(*t* − 250 *ms*), …, *x*_i_(*t*), …, *x*_i_(*t* + 250 *ms*)] where *x*_i_(*t*) represents the inferred firing rates of the *i*^th^ neuron at zero time shift. The ranger function of the ranger R package version 0.10.1 was used to construct regression forests, with the snout movement as the dependent variable and the binned inferred firing rates of a given neuron as predictors. Most arguments of the function were kept at default settings, except the following: the number of trees was set to 128, the minimum size of terminal nodes was set to 2, the number of predictor variables randomly sampled at each node split was set to the maximum between 1 or the third of the number of predictors, and the variable importance mode was set to “impurity”. To obtain a prediction for all trials, 5-fold cross-validation was applied by training the algorithm on 80% of the data (i.e. training set) and evaluating it on the remaining 20% of the data (i.e. test set). For each neuron, the decoding accuracy was assessed by computing the Pearson’s product-moment correlation coefficient between the observed and predicted behavioral event fluctuations.

## Statistical Analysis

All data are expressed as the mean ± s.e.m. unless stated otherwise. No data sets were excluded from analysis. For data obtained from electophysiological recordings, before applying the Student *t*-test, a QQ plot was generated and the Shapiro-Wilk test was performed for each pool of data to confirm a normal distribution. Repeated-measures ANOVA, between-subjects factors ANOVA, mixed-factors ANOVA, and *post-hoc t*-tests were used to test for statistical differences between experimental conditions. Sidak’s correction was used to correct for multiple comparisons. Statistical analyses were performed using Prism (GraphPad), Origin 2021 (OriginLab) or MATLAB and considered significant if *P* < 0.05. Power analyses were performed using G*power and reported as Type II error probability (β).

