## Supplemental information for "Higher-order thalamocortical projections selectively control excitability via NMDAR and mGluRI-mediated mechanisms"

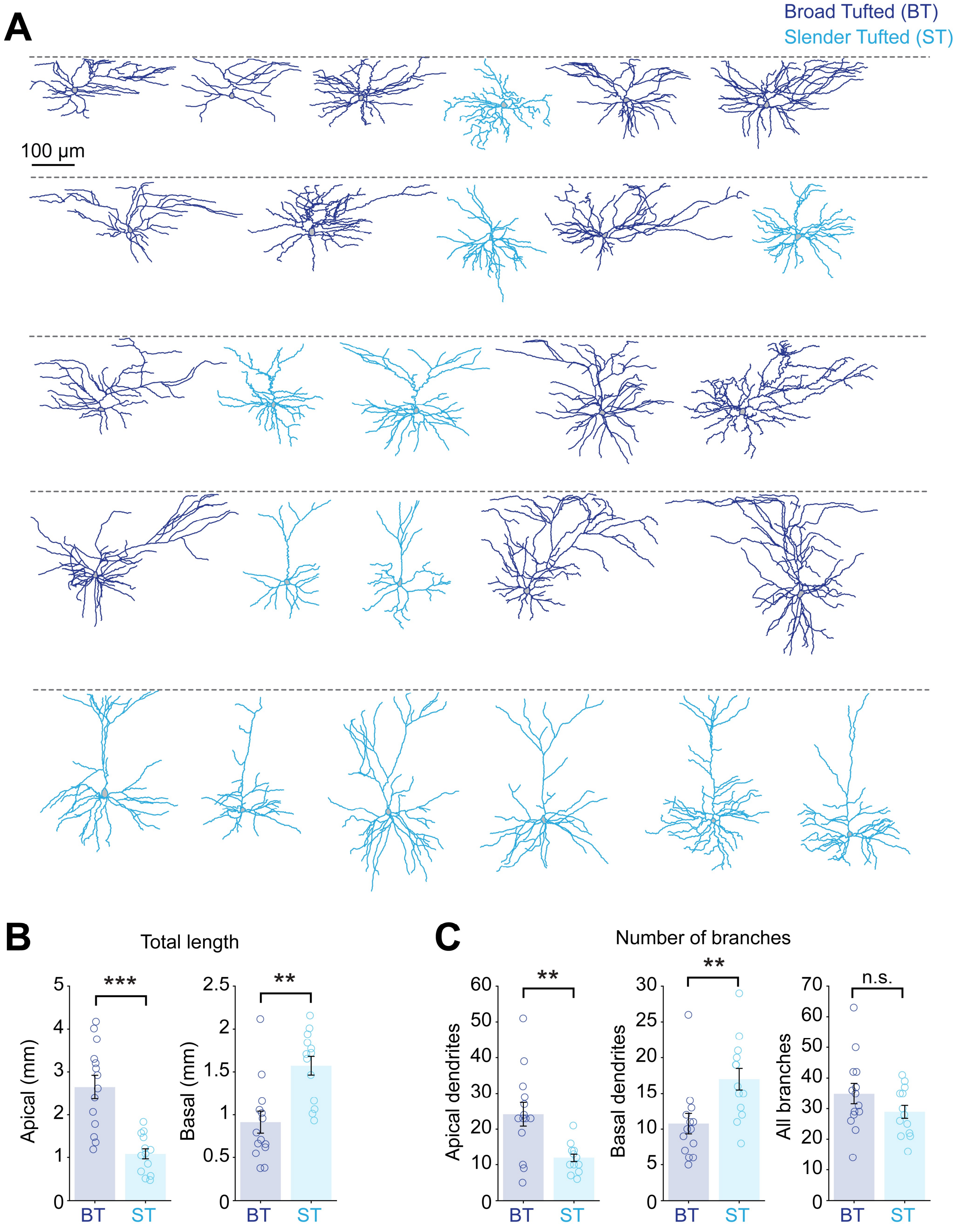


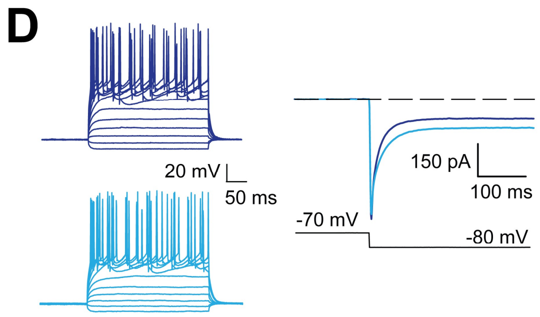


E

| Passive properties | BT (n = 30) | ST (n = 34) | P value |
|---|---|---|---|
| $V_m$ (mV) | -73.20 ± 3.75 | -72.4 ± 2.90 | 0.87 |
| $R_{in}$ (mΩ) | 161.0 ± 16.0 | 177.4 ± 18.0 | 0.38 |
| $\tau_{fast}$ (ms) | 0.43 ± 0.06 | 0.68 ± 0.13 | 0.08 |
| $\tau_{slow}$ (ms) | 2.2 ± 0.2 | 4.2 ± 0.8 | < 0.02 * |
| Capacitance (pF) | 43.1 ± 2.56 | 78.4 ± 4.7 | < 0.01 ** |

**Figure S1. L2/3 pyramidal neurons in the S1 can be segregated into two groups based on their morphology and electrophysiological features (related to Figure 1)**

**A**- Morphological reconstructions of the dendritic arborization of 27 recorded and biocytin-filled L2/3 pyramidal neurons. Neurons are sorted ascendingly by the depth of their soma relative to the pia (represented by a dotted line). BT and ST neurons were segregated based on the clustering analysis of the dendritic span and density within the first 200 µm (cf. Figure 1, BT neurons are colored in dark blue and ST neuron are colored in light blue). **B**- BT neurons have longer apical dendrites than ST neurons ($n$ = 14 BT and 13 ST neurons; $P$ = 1.7×$10^{-4}$, Wilcoxon ranked sum test) but have shorter basal dendrites ($P$ = 0.0015, Wilcoxon ranked sum test). **C**- BT neurons have more apical branches as compared to ST neurons ($n$ = 14 BT and 13 ST neurons; $P$ = 0.005, Wilcoxon ranked sum test) but have less basal dendrites ($P$ = 0.003, Wilcoxon ranked sum test). On the right, the comparison of the total number of branches in not significantly different between BT and ST neurons ($P$ = 0.17, Wilcoxon ranked sum test). **D**- Examples traces of the firing pattern of a BT (top) and a ST neuron (middle) in response to a series of current steps displaying very similar firing properties. The passive properties were assessed by analyzing the changes in current to a hyperpolarizing step from -70 mV to -80 mV (bottom). **E**- Summary table of the passive properties of BT and ST neurons (Wilcoxon signed-rank test used for all parameters). Data displayed as mean ± s.e.m.

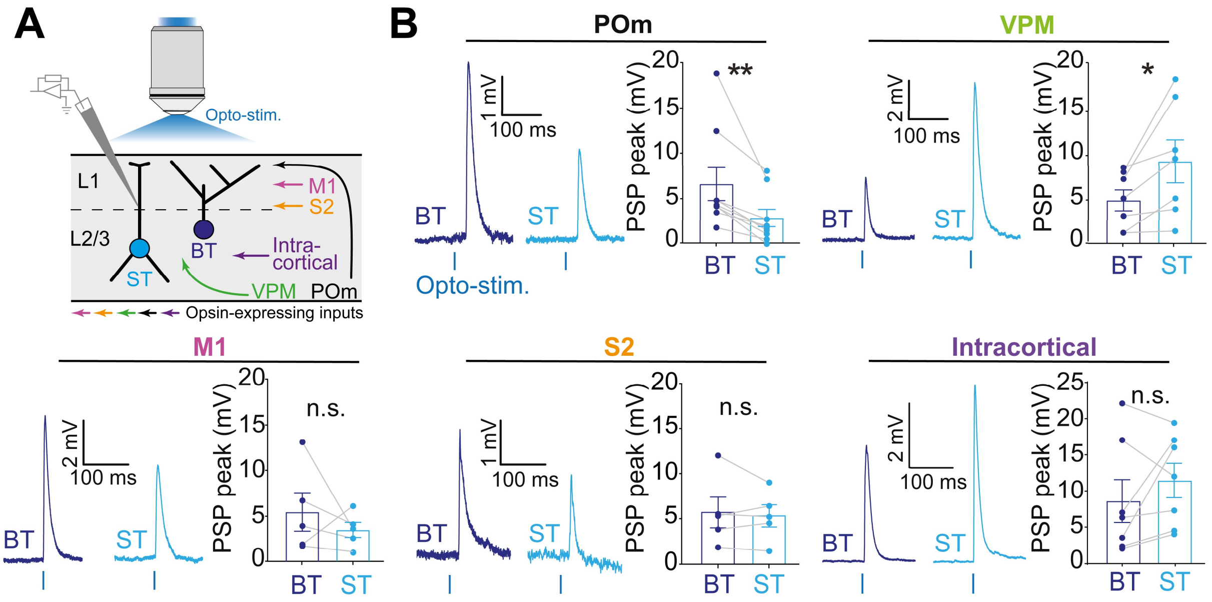


**Figure S2. Data pairing of BT and ST neurons within brain slices to account for the variability of opsin expression of the various inputs (related to Figure 2)**

**A**- Example of an acute brain slice where a BT and a ST neuron were recorded and biocytin-filled. To compare the relative weight of the various input onto BT and ST neurons by accounting for the variability of the opsin expression, dendritic recordings were performed on both cell types in each brain slices. **B**- YFP signal reflecting the ChR2 expression in the thalamocortical projections from POm in the same brain slice shown in A (left). To control for the homogenous expression of the opsin in L1, a series of regions of interest (squares of 50 µm) were drawn below the pia to measure the average fluorescence of the YFP signal. In this example, the quantification shows that for these neighboring recorded cells, the expression is relatively homogenous (right). Scale bars are 100 µm. **C**- PSP response amplitudes evoked by POm photostimulation of ST and BT dendritic recordings as a function of the average YFP expression reveals a linear correlation between YFP and ChR2 expression (for BT neurons, $R^2 = 0.83$; for ST neurons, $R^2 = 0.73$, $n = 11$ pairs of neurons and slices). **D**- To control that the differences in amplitude observed between BT and ST neurons were not explained by the dampening of the PSPs due to different electronic distances, PSP rise times were compared for all measured inputs. The rise time of a PSP was defined as the time difference between the 10% to the 90% of the maximum value of the PSP. No difference in rise time between BT and ST neurons was observed for all the tested inputs (for POm, BT: 2.1 ± 0.2 ms, $n = 9$, ST: 2.3 ± 0.3 ms, $n = 9$, $P = 0.64$; for VPM, BT: 3.9 ± 0.6 ms, $n = 7$, ST: 3.7 ± 0.4 ms, $n = 7$, $P = 0.75$; for M1, BT: 2.7 ± 0.1 ms, $n = 5$, ST: 2.5 ± 0.2 ms, $n = 5$, $P = 0.58$; for S2, BT: 3.0 ± 0.4 ms, $n = 5$, ST: 2.9 ± 0.4 ms, $n = 5$, $P = 1$; for $S1_{intracortical}$, BT: 2.6 ± 0.2 ms, $n = 7$, ST: 2.7 ± 0.2 ms, $n = 7$, $P = 0.78$, Wilcoxon signed-rank tests).

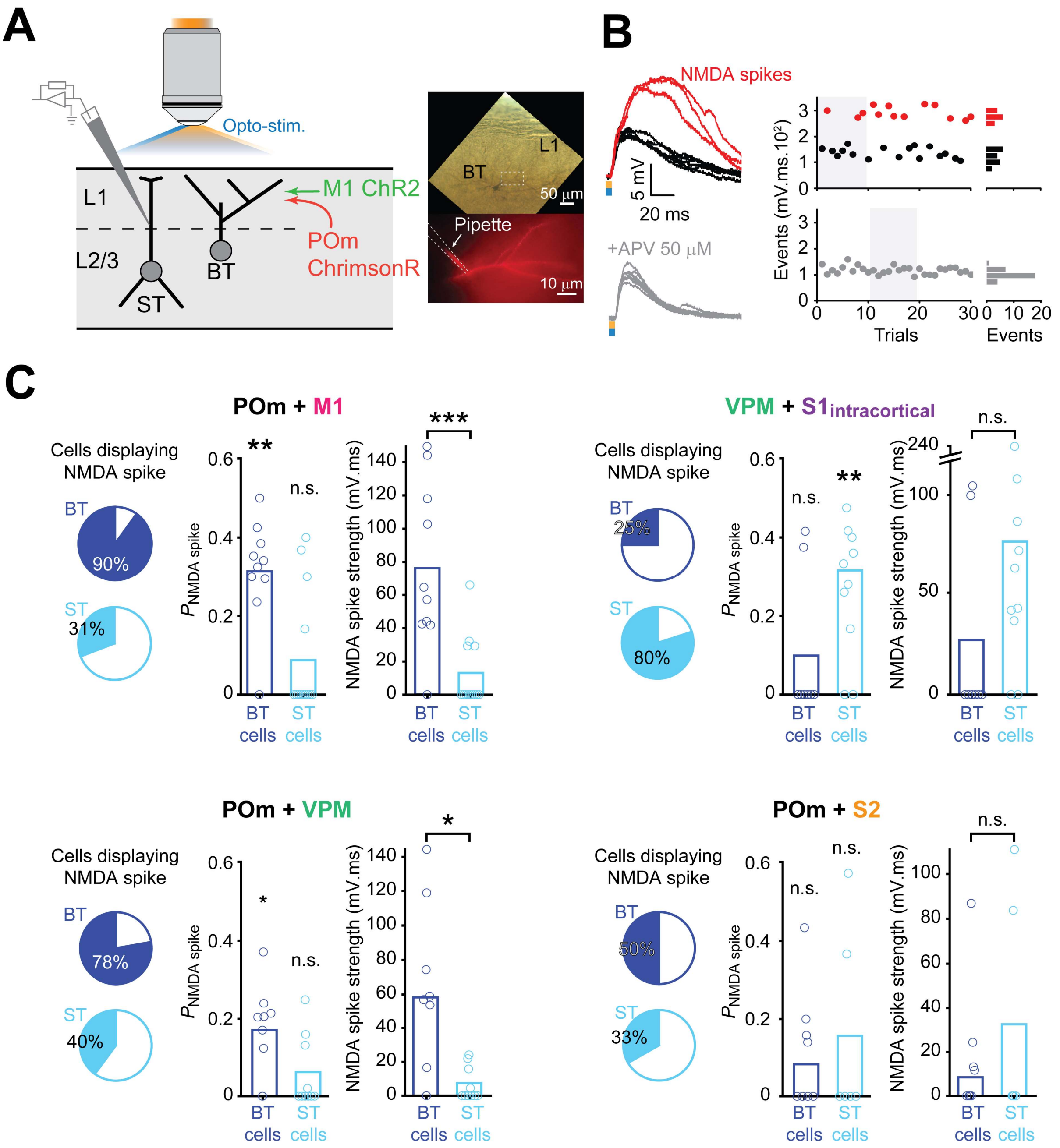


**Figure S3. Comparison of the monosynaptic response of POm and VPM in BT and ST neurons (related to Figure 2)**

**A**- Dendritic recordings in a BT neuron (left) and a ST neuron (right) showing the evoked PSPs when opsins-expressing POm and VPM inputs are photostimulated independently (5 pulses of 1 ms at 8 Hz) before and after bath applying TTX (1 µM) and 4-AP (100 µM) to reveal the direct monosynaptic response. The responses to the POm stimulations were reduced for both neuron types. However, the responses to the VPM stimulations were completely abolished in the BT neuron. **B**- Comparison of the mean amplitude of the PSP responses to POm or VPM stimulations for BT and ST neurons in control versus in TTX + 4-AP conditions. Notably, the application of TTX + 4-AP fully abolished the response to VPM stimulation in BT neurons. The noise level of the recording was determined as the minimum and maximum values of the pre-stimulation baseline period.

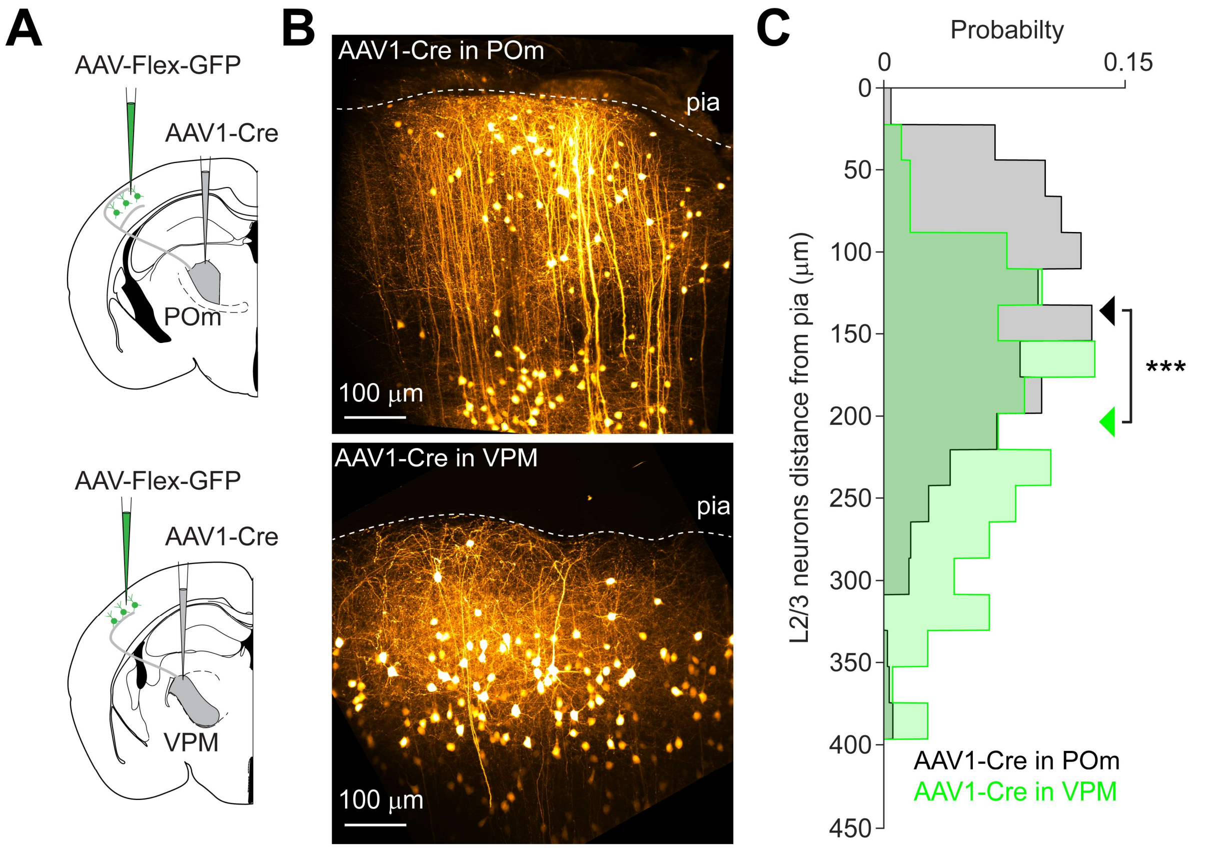


**Figure S4. Trans-synaptic AAV delivery from POm and VPM to S1 neurons (related to Figures 1 and 2)**

**A**- An AAV1 vector expressing Cre recombinase was injected locally into either the POm or VPM. This allowed for anterograde trans-synaptic expression of Cre recombinase in downstream neurons, including those in the S1 barrel cortex. Subsequently, a second AAV vector expressing a Cre-dependent conditional GFP reporter was injected into the S1 barrel cortex, enabling selective expression of GFP in the subset of neurons that received input from the targeted thalamic neurons expressing Cre recombinase. **B**- Standard deviation projections of 2-photon image stacks of the GFP-expressing neurons in S1 when the AAV1-Cre was injected in the POm (top) and in the VPM (bottom). Scale bars are 100 µm. **C**- Distribution of L2/3 neurons' somata depth expressing GFP. The anterograde trans-synaptic labeling from the POm targeted more superficial L2/3 neurons than the VPM (For POm: mean ± s.d., 136 ± 70 µm, *n* = 897 neurons from 5 mice; For VPM: mean ± s.d., 204 ± 79 µm, *n* = 183 neurons from 2 mice, $P = 4.52\times10^{-17}$, Kolmogorov-Smirnov test).

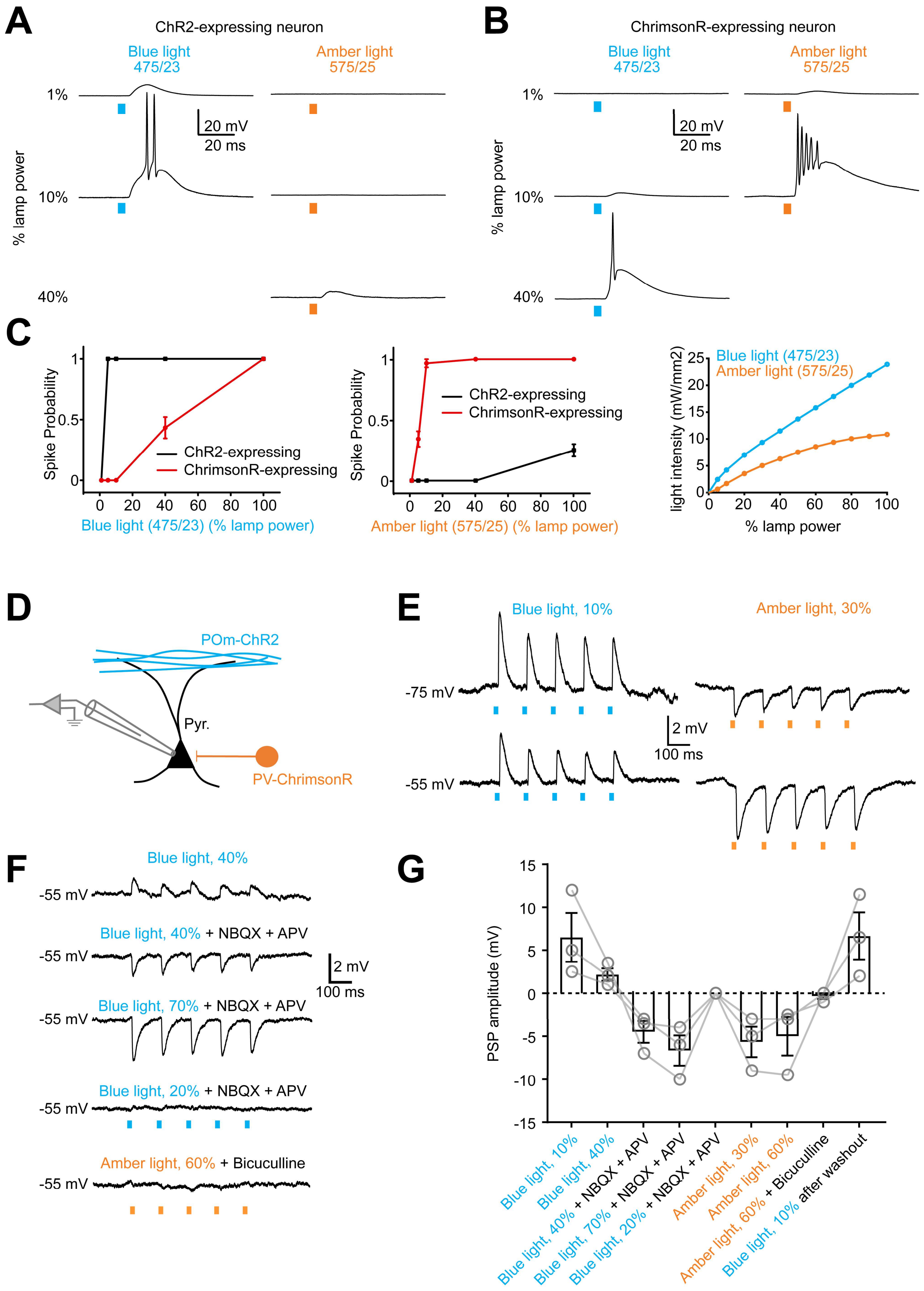
A
ChR2-expressing neuron
Blue light
475/23
Amber light
575/25
1%
10%
40%
% lamp power
20 mV
20 ms
B
ChrimsonR-expressing neuron
Blue light
475/23
Amber light
575/25
1%
10%
40%
% lamp power
20 mV
20 ms
C
Spike Probability
ChR2-expressing
ChrimsonR-expressing
Blue light (475/23) (% lamp power)
Amber light (575/25) (% lamp power)
light intensity (mW/mm2)
Blue light (475/23)
Amber light (575/25)
% lamp power
D
POm-ChR2
Pyr.
PV-ChrimsonR
E
Blue light, 10%
Amber light, 30%
-75 mV
-55 mV
2 mV
100 ms
F
Blue light, 40%
Blue light, 40% + NBQX + APV
Blue light, 70% + NBQX + APV
Blue light, 20% + NBQX + APV
Amber light, 60% + Bicuculline
-55 mV
2 mV
100 ms
G
PSP amplitude (mV)
Blue light, 10%
Blue light, 40%
Blue light, 40% + NBQX + APV
Blue light, 70% + NBQX + APV
Blue light, 20% + NBQX + APV
Amber light, 30%
Amber light, 60%
Amber light, 60% + Bicuculline
Blue light, 10% after washout

1475

**Figure S5. Calibration of the light dose in the double-opsins experiments (related to Figure 3)**

**A**- Somatic patch-clamp recordings and photostimulation of a ChR2-expressing neuron with a 10-ms pulse of blue light (475/23 nm) or amber light (575/25 nm) at various intensities. The neuron spiked when 10% of the blue light power is used whereas subthreshold activity is evoked with 40% of amber light power. **B**- Same experiment with a neuron expressing ChrimsonR. In this case, 10% of amber light is sufficient to elicit spiking of the neuron but strong blue light power (40%) may induce the spiking of the neuron as well. **C**- Left, spike probability as a function of blue light power for ChR2- and ChrimsonR-expressing neurons (black and red lines, respectively). Using blue light power at 20% reliably made the ChR2-expressing neurons spike with minimal stimulation of the ChrimsonR-expressing neurons. Center, spike probability as a function of amber light power for ChR2- and ChrimsonR-expressing neurons. Again, a lamp power of 20% maximized the spike probability of ChrimsonR-expressing neurons without exciting ChR2-expressing neurons ($n = 3$ neurons, probability calculated from 20 trials for each opsin and each power intensity). On the right, correspondence of the blue and amber light intensity as a function of the lamp power expressed in percentage. **D**- Test of the light-evoked synaptic responses in L2/3 pyramidal neurons from independent excitation of two different inputs. Specific expression of ChrimsonR in PV interneurons was performed by injecting a Cre-dependent AAV vector in S1 of a PV-Cre transgenic mouse, and a local injection of a non-Cre dependent AAV vector was used to express ChR2 in the POm. **E**- Examples of current-clamp recordings of L2/3 pyramidal neuron responses to blue or amber light stimulation with a train of 5 pulses of 1 ms at 8 Hz. At resting membrane potential, 10% blue light elicits excitatory PSPs (EPSPs) while 30% amber light elicits inhibitory PSPs (IPSPs). Holding the membrane potential at -55 mV (as it was done for most of the experiments in this study) only affects the amplitudes of the responses due to a change in the driving force but not their sign. **F**- Increasing the blue light power up to 40% decreases the EPSP amplitude due to the simultaneous activation of ChrimsonR in PVs. Blocking the glutamatergic response using 10 µM NBQX and 50 µm APV reveals the inhibitory component of the response, confirming the contamination of the response from ChrimsonR activation at this power. Increasing the blue light power up to 70% amplifies the IPSP response while 20% of blue light power does not elicit any IPSP. In another condition, increasing the power up to 60% when the GABAergic response is blocked with 10 µM bicuculline does elicit any EPSP. **G**- Averaged PSP amplitude obtained with various light stimulations and bath applied drugs. The data indicates that ChrimsonR is not activated when the blue light power is below 20% and ChR2 is not activated when the amber light power is below 60% of the maximum lamp power ($n = 3$ neurons).

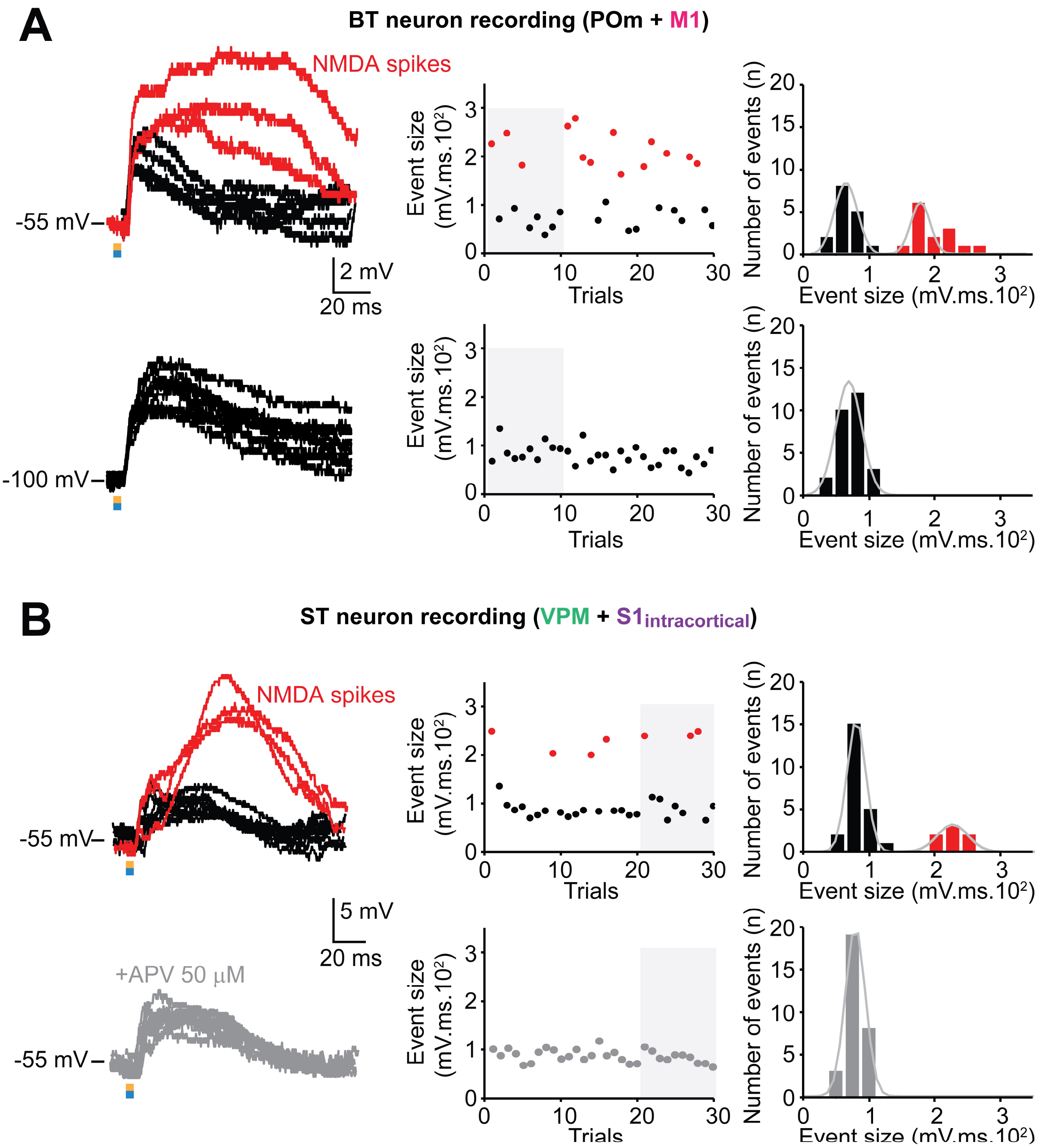


**Figure S6. NMDA spikes are prevented in hyperpolarized conditions and are also NMDAR-dependent in ST neurons (related to Figure 3)**

**A**- Dendritic recordings of a BT neuron in S1 during the co-stimulation of POm and M1 inputs. NMDA spikes were observed when the cell was slightly depolarized at -55 mV (top) but were prevented when the cell was hyperpolarized at -100 mV (bottom). While NMDA spikes are prevented at -100 mV, regular PSP event size was not different from the condition at -55 mV.

**B**- Dendritic recordings of a ST neuron in S1 during the co-stimulation of VPM and $S1_{intracortical}$ inputs also displayed NMDA spikes (top). Bath application of APV (50 µM) prevented the occurrence of NMDA spikes (bottom).

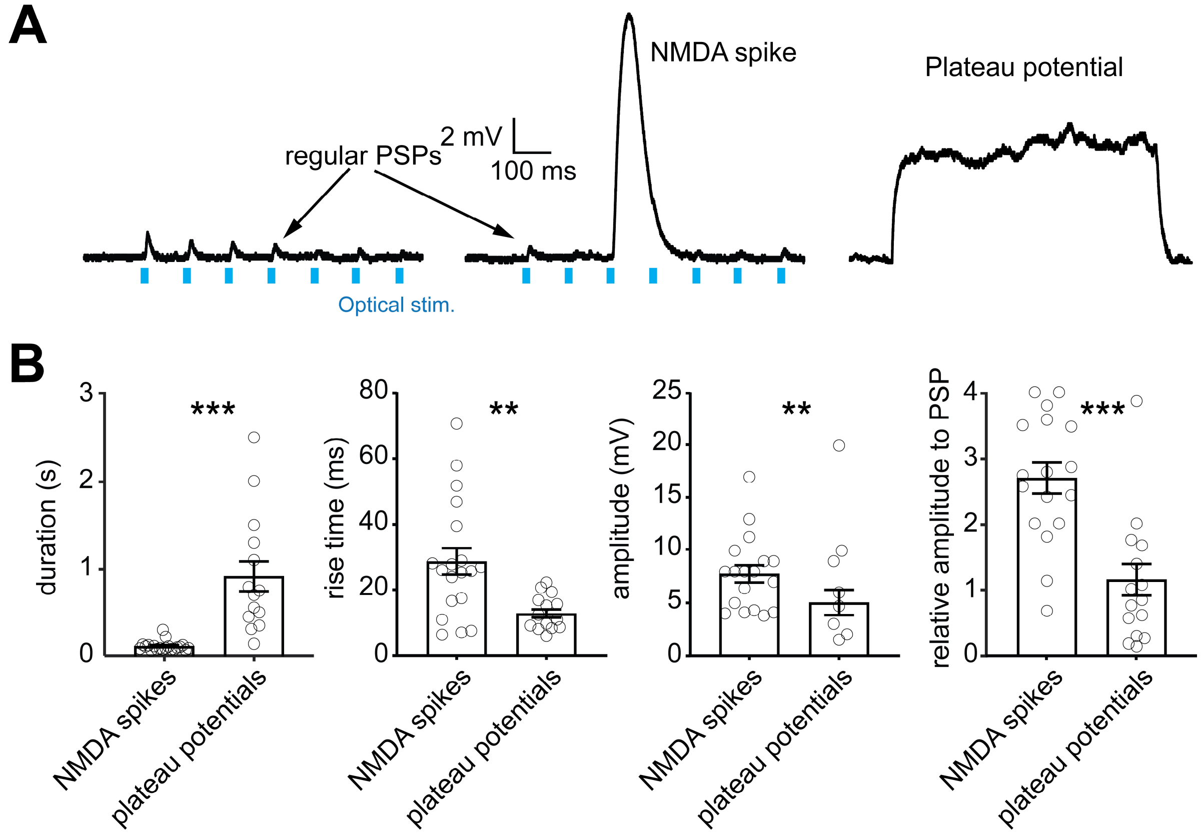


**Figure S7. Comparison between NMDA spikes and plateau potentials (related to Figure 4)**

**A**- Examples of dendritic recordings of a BT neuron during simultaneous photostimulation of POm and M1 afferent inputs. Evoked PSPs, NMDA spikes and delayed plateau potentials were observed in this recording. **B**- Comparison of the duration, rise time, amplitude and relative amplitude to PSP between NMDA spikes and plateau potentials. Plateau potentials are characterized by a longer duration (for plateau potentials, 912.9 ± 170.9 ms, $n$ = 16; for NMDA spikes, 112,7 ± 12.3 ms, $n$ = 18, $P < 0.001$, Mann-Whitney test), a faster rise-time (for plateau potentials, 12.7 ± 1.2 ms, $n$ = 16; for NMDA spikes, 28.7 ± 4.0 ms, $n$ = 18, $P$ = 0.0018, Mann-Whitney test), a smaller amplitude (for plateau potentials, 5.0 ± 1.2 mV, $n$ = 16; for NMDA spikes, 7.7 ± 0.8 mV, $n$ = 18, $P$ = 0.0079, Wilcoxon test), and a smaller relative amplitude to PSP (for plateau potentials, 1.1 ± 0.2, $n$ = 16; for NMDA spikes, 2.7 ± 0.2, $n$ = 18, $P <$ 0.0001, Mann-Whitney test).

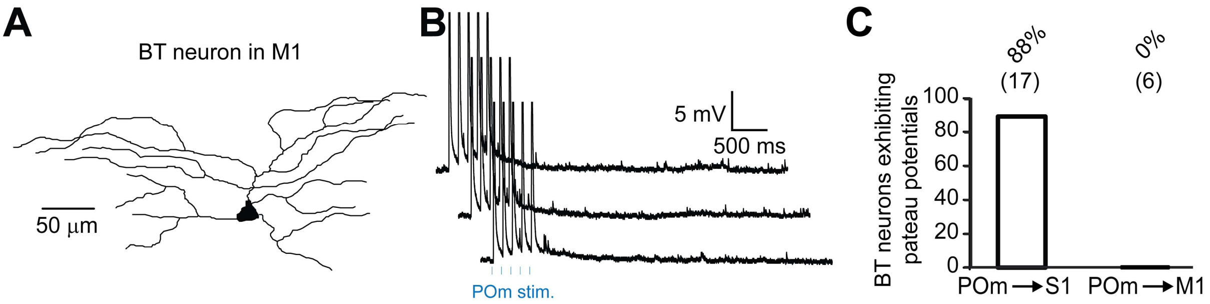


**Figure S8. BT neurons in M1 do not exhibit plateau potentials (related to Figure 4)**
**A**- Example of the morphological reconstruction of a BT neuron located in M1. **B**- Dendritic recordings of a BT neuron in M1 during and after the optogenetic stimulation of POm. Unlike for BT neurons in S1, no plateau potentials were observed following the stimulation of POm. **C**- Fraction of BT neurons exhibiting plateau potentials upon POm stimulation in S1 and in M1. None of the cells recorded in M1 displayed a plateau potential (0 out of 6) while 88% of the cells in S1 did (15 out of 17 neurons).

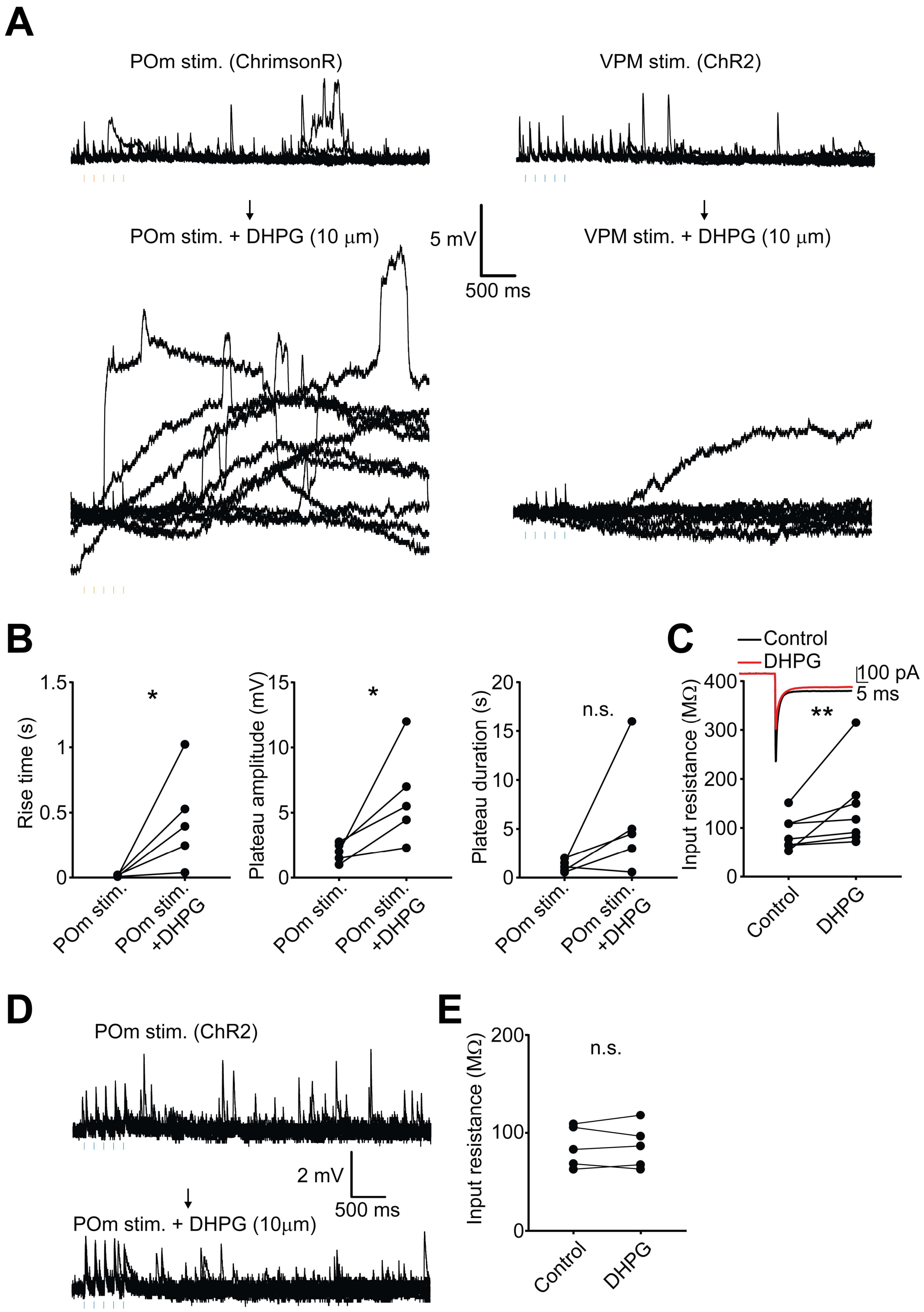


**Figure S9. Effect of DHPG on plateau potentials (related to Figure 6)**

**A**- Example of a dendritic recording of a BT cell after photostimulation of ChrimsonR-expressing POm and ChR2-expressing VPM. mGluRI-mediated plateau potentials were induced only upon POm stimulation. Subsequent bath perfusion of DHPG (10 µM) increased the number, amplitude, and duration of events after the photostimulation of POm afferent inputs. However, this did not increase the occurrence of plateau potentials after VPM stimulation, indicating that DHPG alone is not sufficient to induce them. **B**- The rise time of plateau potentials, following POm photostimulation, significantly increased in the presence of DHPG (control: 13.7 ± 3.1 ms; DHPG: 446.2 ± 165.5 ms, $n = 5$, $P = 0.05$, paired t-test). It also significantly increased the amplitude of these events (control: 1.9 ± 0.3 mV; DHPG: 6.2 ± 1.6 mV, $n = 5$, $P = 0.05$, paired t-test) but not their duration (control: 1180.9 ± 260.5 ms; DHPG: 5813.3 ± 2660.8 ms, $n = 5$, $P = 0.14$, paired t-test). **C**- Perfusion of DHPG significantly increased the dendritic input resistance of BT neurons (control: 79.2 ± 1.2 MΩ; DHPG: 127.2 ± 3.0 MΩ, $n = 8$, $P = 0.0078$, paired t-test). **D**- Bath application of DHPG in addition to POm stimulation failed to induce plateau potentials in ST neurons. **E**- DHPG did not change the input resistance in ST neurons (control: 85.4 ± 9.4 MΩ; DHPG: 85.6 ± 10.0 MΩ, $n = 5$, $P = 0.88$, paired t-test).

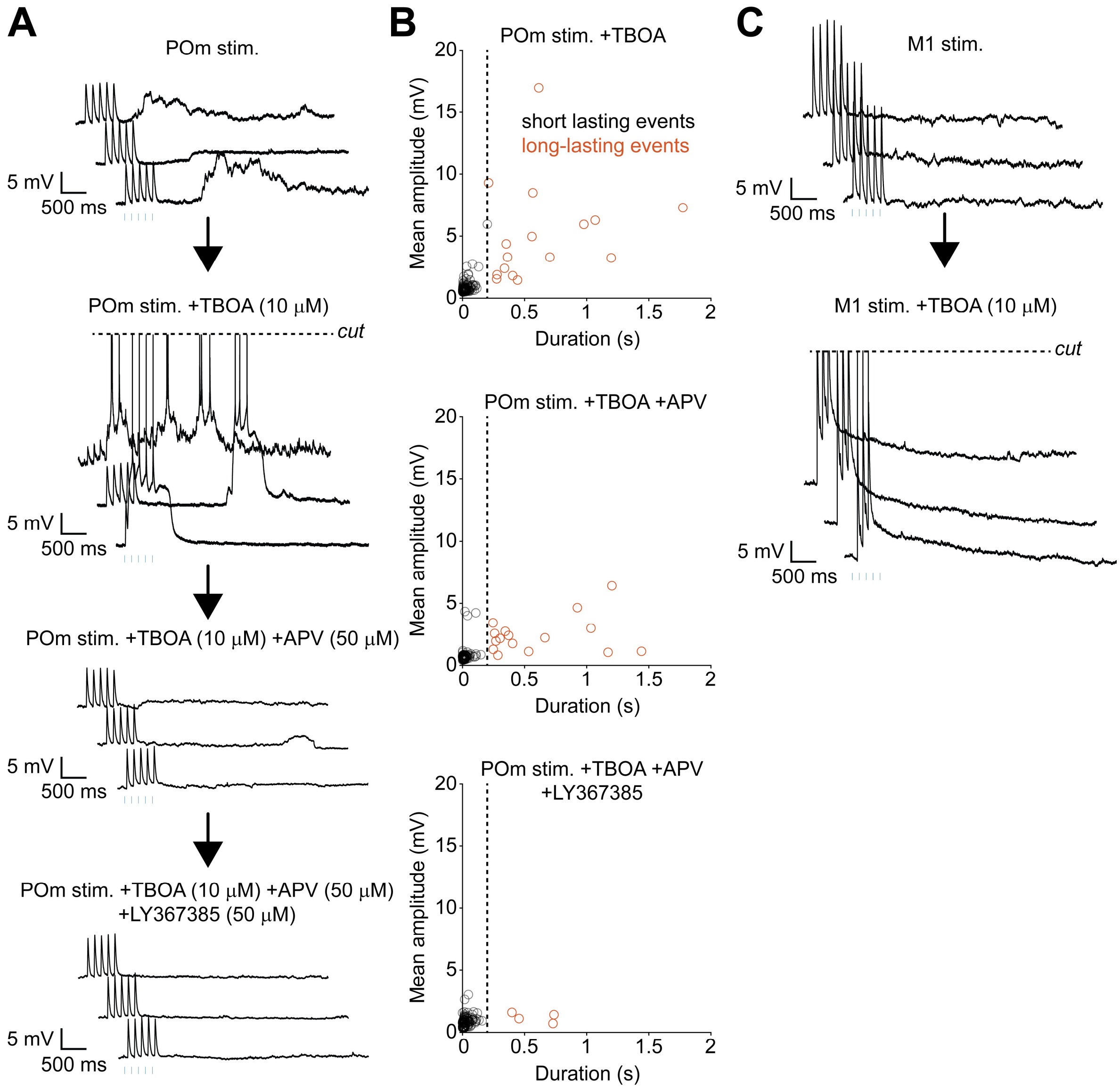


**Figure S10. POm-mediated plateau potentials boost NMDA-mediated spiking when the spontaneous activity is enhanced (related to Figures 6 and 7)**

**A-** Dendritic recordings on a BT neuron in S1 displaying long-lasting and delayed plateau potentials after photostimulation of POm afferent inputs. Bath application of 10 µM TBOA, a glutamate reuptake inhibitor, enhancing the spontaneous activity facilitated the triggering of action potentials during plateau potentials events. Adding 50 µM APV to the bath prevented the generation of these spikes but plateau potentials remained. These were indeed confirmed to be mGluRI-mediated plateau potentials as they were abolished when adding 50 µM LY367385 to the bath. **B**- Scatter plots showing the duration and mean amplitude of all post-stimulation events automatically detected in BT neurons recordings in S1 (*n* = 3). Many large and long-lasting events (> 200 ms), corresponding to action potentials or NMDA spikes and plateau potentials respectively, were detected under TBOA conditions. The addition of APV reduced the number of large events, but long-lasting plateau potentials events remained. Finally, adding LY367385 in the bath resulted in the decrease in number of the detected long-lasting events. **C**- Dendritic recordings on the same neuron shown in (**A**) after photostimulation of M1 afferents inputs did not produce any plateau potentials. M1 photostimulation during bath perfusion of TBOA generated spikes but no delayed spiking events were observed.

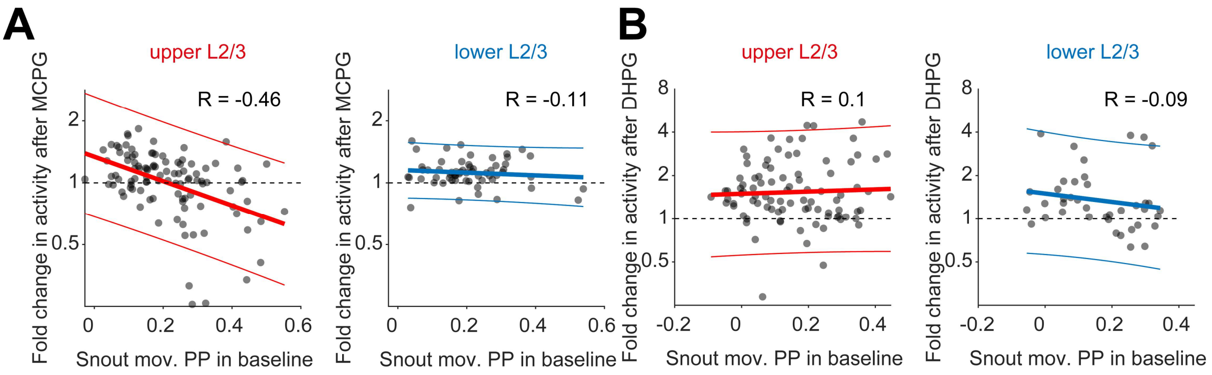


**Figure S11. Relation between movement prediction in baseline and mGluRI modulation of activity (related to Figure 7)**

**A**- Correlation between the snout movement PP in baseline and the change in activity after MCPG injection. Upper L2/3 neurons that show the strongest reduction in activity in MCPG predicted best the snout movement during the baseline recording ($n$ = 104 upper and 61 lower L2/3 neurons, from 3 mice; Pearson's correlation $R$ = -0.46). Lower L2/3 neurons show little to no correlation ($R$ = -0.11). **B**- Same for DHPG dataset, little to no correlation were observed for the upper and lower L2/3 neurons ($n$ = 90 upper and 40 lower L2/3 neurons, from 5 mice; R= 0.1 and -0.09 respectively).
